# Differential N-end rule degradation of RIN4/NOI fragments generated by the AvrRpt2 effector protease

**DOI:** 10.1101/583054

**Authors:** Kevin Goslin, Lennart Eschen-Lippold, Christin Naumann, Eric Linster, Maud Sorel, Maria Klecker, Rémi de Marchi, Anne Kind, Markus Wirtz, Justin Lee, Nico Dissmeyer, Emmanuelle Graciet

## Abstract

The protein RPM1-INTERACTING PROTEIN4 (RIN4) is a central regulator of both layers of plant immunity systems, the so-called pattern-triggered immunity (PTI) and effector-triggered immunity (ETI). RIN4 is targeted by several effectors, including the *Pseudomonas syringae* protease effector AvrRpt2. Cleavage of RIN4 by AvrRpt2 generates unstable RIN4 fragments, whose degradation leads to the activation of the resistance protein RPS2 (RESISTANT TO P. SYRINGAE2). Hence, identifying the determinants of RIN4 degradation is key to understanding RPS2-mediated ETI, as well as virulence functions of AvrRpt2. In addition to RIN4, AvrRpt2 cleaves host proteins from the nitrate-induced (NOI) domain family. Although cleavage of NOI-domain proteins by AvrRpt2 may contribute to PTI regulation, the (in)stability of these proteolytic fragments and the determinants that regulate their stability have not been examined. Notably, a common feature of RIN4 and of many NOI-domain protein fragments generated by AvrRpt2 cleavage is the exposure of a new N-terminal residue that is destabilizing according to the N-end rule. Using antibodies raised against endogenous RIN4, we show that the destabilization of AvrRpt2-cleaved RIN4 fragments is independent of the N-end rule pathway (recently renamed N-degron pathway). By contrast, several NOI-domain protein fragments are *bona fide* substrates of the N-degron pathway. The discovery of this novel set of substrates considerably expands the number of proteins targeted for degradation by this ubiquitin-dependent pathway, for which very few physiological substrates are known in plants. Our results also open new avenues of research to understand the role of AvrRpt2 in promoting bacterial virulence.

**One sentence summary:** Analysis of RIN4/NOI fragments released after cleavage by the bacterial effector protease AvrRpt2 reveals a novel role of the N-end rule in the degradation of NOI-domain proteins, but not of RIN4.

## Introduction

Plants have evolved complex mechanisms to fight off pathogens. A first line of defense is initiated through the recognition of pathogen-associated molecular patterns (PAMPs) by surface-localized transmembrane pattern recognition receptors (PRRs), resulting in the activation of multiple signal transduction pathways, large transcriptional changes and the onset of pattern-triggered immunity (PTI) (Jones and Dangl, 2006; Henry et al., 2013; Couto and Zipfel, 2016). Pathogens also code for effector proteins or molecules that are secreted. These effectors mis-regulate different aspects of the PTI response or upstream signalling cascades by hijacking or manipulating the function of host proteins. In the absence of cognate receptors for these effectors, their activity results in dampened host immunity and increased pathogen virulence. However, these effectors may be detected, directly or indirectly, by intracellular nucleotide binding site leucine rich repeat receptor proteins (NBS-LRR). This recognition elicits a stronger response termed effector-triggered immunity (ETI), which is often associated with a localized programmed cell death (Jones and Dangl, 2006; van der Hoorn and Kamoun, 2008; Kourelis and van der Hoorn, 2018).

A key regulator of plant immunity is the membrane-bound protein RPM1-INTERACTING PROTEIN4 (RIN4), which acts as a negative regulator of both PTI and ETI (Day et al., 2005; Kim et al., 2005b; Liu et al., 2009; Afzal et al., 2011; Toruno et al., 2019). Notably, RIN4 is targeted by multiple effector proteins, including AvrRpm1 (Mackey et al., 2002), AvrB (Mackey et al., 2002; Desveaux et al., 2007), HopF2 (Wilton et al., 2010) and HopZ3 (Lee et al., 2015b). The effector protease AvrRpt2 also targets RIN4 (Axtell et al., 2003; Mackey et al., 2003; Chisholm et al., 2005), as well as other proteins that have the AvrRpt2 consensus recognition sequence PxFGxW (Chisholm et al., 2005; Kim et al., 2005a; Eschen-Lippold et al., 2016a). Following delivery into host cells and plant cyclophilin-dependent activation (Jin et al., 2003; Coaker et al., 2005), AvrRpt2 undergoes autocatalytic cleavage (Axtell et al., 2003; Chisholm et al., 2005) and cleaves RIN4 at two specific sites within the N-terminal or C-terminal nitrate-induced (NOI) domains of RIN4. These are referred to as RCS1 and RCS2 (RIN4 cleavage site 1 or 2), respectively (Fig. 1A). In Arabidopsis *rpm1 rps2* double mutant plants lacking functional RESISTANCE TO P. SYRINGAE PV MACULICOLA1 (RPM1) and RESISTANT TO P. SYRINGAE2 (RPS2) NBS-LRR proteins, these RIN4 fragments supress PTI (Afzal et al., 2011). RIN4 and its cleavage by AvrRpt2 may also play a role in the regulation of EXO70B1, a subunit of the exocyst complex that is important for autophagic-related protein trafficking (Kulich et al., 2013; Sabol et al., 2017) and plays a role in plant immunity (Stegmann et al., 2013; Liu et al., 2017). Notably, AvrRpt2 also promotes virulence through RIN4-independent mechanisms, including the manipulation of auxin signalling (Chen et al., 2007; Cui et al., 2013) and the repression of mitogen-activated protein kinase (MAPK) pathways (Eschen-Lippold et al., 2016a; Eschen-Lippold et al., 2016b).

**Figure 1:**
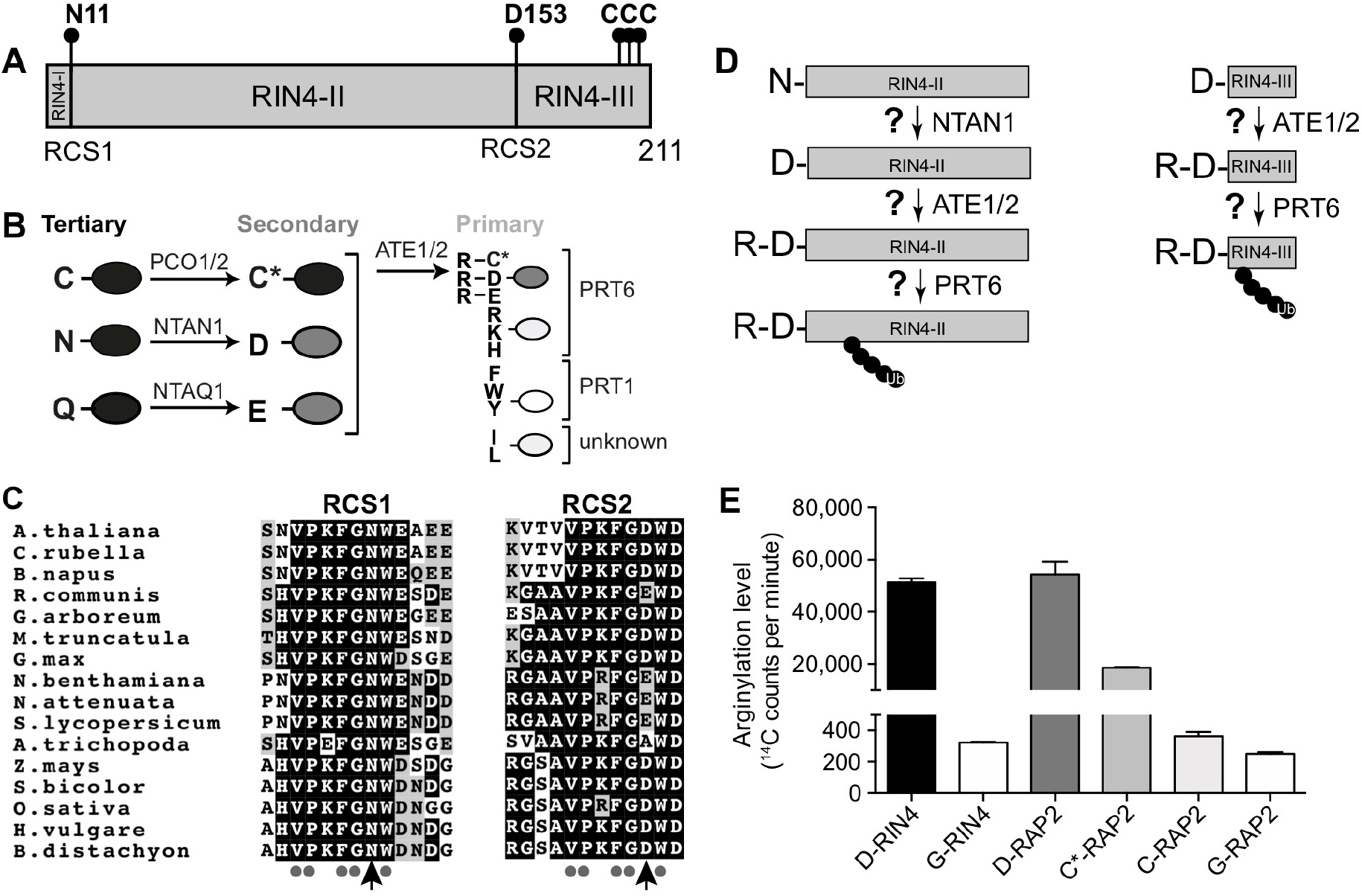
AvrRpt2 cleavage sites and neo-N-terminal residues of RIN4 are conserved and can act as putative N-degron. **(A)** Scheme of AvrRpt2 cleavage sites in Arabidopsis RIN4. RCS1: RIN4 cleavage site 1; RCS2: RIN4 cleavage site 2; RIN4-I: N-terminal fragment of RIN4 released after cleavage at RCS1; RIN4-II: RIN4 fragment following AvrRpt2 cleavage at both RCS1 and RCS2; RIN4-III: C-terminal fragment of RIN4 released after cleavage at RCS2 by AvrRpt2. Newly exposed N-terminal residues (N11 at the N-terminus of RIN4-II and D153 at the N-terminus of RIN4-III are indicated). The three C-terminal cysteine residues (positions 203 to 205) that are palmitoylated and serve to target RIN4 to the plasma membrane are also indicated. **(B)** Hierarchical organization of the Arg/N-degron pathway in Arabidopsis. Destabilizing N-terminal residues may target proteins for degradation by the ubiquitin-dependent N-degron pathway. Primary destabilizing N-terminal residues are directly recognized by E3 ligases (or N-recognins) of the N-degron pathway, including PRT6 and PRT1. In contrast, secondary destabilizing residues are first modified by Arg-transferases that conjugate Arg at the N-terminus of proteins starting with Asp, Glu and oxidized Cys (noted Cys*). In addition, tertiary destabilizing residues are first either oxidized by PLANT CYSTEINE OXIDASE (PCO) enzymes (in the case of Cys) or deamidated into Asp and Glu by Asn- and Gln-specific deamidases, NTAN1 and NTAQ1, respectively. **(C)** The RCS1 and RCS2 cleavage sites are evolutionarily conserved in plants. Sequence alignment of RCS1 and RCS2 sites from different RIN4 orthologs identified using NCBI BLASTp with Arabidopsis RIN4 (At3g25070) as a query. *A. thaliana*: At3g25070, corresponding to NP_189143.2; *C. rubella*: *Capsella rubella* XP_023641759.1; *B. napus*: *Brassica napus* XP_013674753.1; *R. communis*: *Ricinus communis* XP_002532749.2; *G. arboreum*: *Gossypium arboreum* KHG28908.1; *M. truncatula*: *Medicago truncatula* XP_013444158.1; *G. max*: *Glycine max* NP_001239973.1; *S. lycopersicum*: *Solanum lycopersicum* XP_010326284.1; *N. benthamiana*: *Nicotiana benthamiana* APY20266.1; *N. attenuata*: *Nicotiana attenuata* XP_019249122.1; *A. trichopoda*: *Amborella trichopoda* XP_011629172.1; *S. bicolor*: *Sorghum bicolor* XP_002444713.2; *Z. mays*: *Zea mays* ONM03164.1; *H. vulgare*: *Hordeum vulgare* AEV12220.1; *B. distachyon*: *Brachypodium distachyon* XP_003572426.1; *O. sativa*: *Oryza sativa* BAF24212.1. **(D)** Schematic representation of the different enzymatic modifications required for N-degron-mediated degradation of the RIN4-II and RIN4-III fragments. **(E)** *In vitro* arginylation of 12-mer peptides derived from the RIN4-II fragment N-terminal region and from known Arg-transferase substrates. RIN4-II peptides with either N-terminal Asp or Gly (noted D-RIN4-II and G-RIN4-II, respectively; X-WEAEENVPYTA) were synthesized and used in *in vitro* arginylation assays. Peptides corresponding to the conserved N-terminal sequence of the RAP2.2 and RAP2.12 proteins, which are known Arg-transferase substrates were also synthesized. Different variants of the latter were used in the *in vitro* arginylation assays, including peptides with N-terminal Asp, oxidized Cys (denoted C*) and Gly (peptides noted D-RAP2, C*-RAP2 and G-RAP2; X-GGAIISDFIPP). Data shown are the average of 2 independent replicates. Error bars represent standard deviations. C* denotes tri-oxidized cysteine (sulfonic acid, R-SO3H).

In an RPS2 genetic background, cleavage of RIN4 at RCS1 and RCS2 triggers RPS2-mediated ETI (Axtell and Staskawicz, 2003; Mackey et al., 2003; Chisholm et al., 2005; Kim et al., 2005a). The RIN4-II and RIN4-III fragments (Fig. 1A) resulting from cleavage at RCS1 and RCS2, respectively, were shown to be unstable (Axtell et al., 2003; Axtell and Staskawicz, 2003; Mackey et al., 2003), although in some cases, they may be detected for up to 6 hrs after AvrRpt2 expression or delivery to host cells (Kim et al., 2005a; Afzal et al., 2011). It has nevertheless been proposed that the degradation of these RIN4 fragments could play a role in the activation of RPS2-mediated ETI (Axtell et al., 2003; Axtell and Staskawicz, 2003; Mackey et al., 2003). Experiments with short N-terminal fragments of RIN4 fused to the N-terminus of GFP suggested that the ubiquitin-dependent N-end rule pathway, which targets proteins for degradation based on the nature of a protein’s N-terminal residue (or N-degron), may play a role in the degradation of the RIN4 fragments (Takemoto and Jones, 2005). However, it is known that recognition of a substrate protein by components of the N-end rule pathway (renamed ‘N-degron pathway’ to reflect recent advances (Varshavsky, 2019)) depends additionally on the conformation of the protein and the accessibility of the N-terminal residue. Hence, a question that has remained unanswered is whether the N-degron pathway can indeed target for degradation the native RIN4-II and III fragments.

While several branches of the N-degron pathways have been uncovered and dissected in yeast and mammals (Varshavsky, 2019), in plants, the so-called Arg/N-degron branch has been the most extensively studied (reviewed in (Gibbs et al., 2014; Gibbs et al., 2016; Dissmeyer et al., 2018; Dissmeyer, 2019)). When present at the N-terminus, so-called primary destabilizing residues of the Arg/N-degron pathway can be recognized by specific E3 ubiquitin ligases termed N-recognins. In Arabidopsis, at least two N-recognins with different substrate specificities exist, namely PROTEOLYSIS1 (PRT1) and PRT6 (Potuschak et al., 1998; Stary et al., 2003; Garzon et al., 2007). Other N-terminal residues function as secondary destabilizing residues and require arginylation (i.e. conjugation of Arg, a primary destabilizing residue) by Arg-transferases (ATE1 and ATE2 in Arabidopsis) before they are recognized by PRT6. Lastly, proteins starting with tertiary destabilizing residues need to be modified before they are arginylated by ATE1/ATE2 and targeted for degradation by PRT6 (Fig. 1B). The plant Arg/N-degron pathway has been shown to regulate various developmental and physiological processes in Arabidopsis (Graciet et al., 2009; Holman et al., 2009; Abbas et al., 2015; Gibbs et al., 2018; Zhang et al., 2018) and in barley (Mendiondo et al., 2016; Vicente et al., 2017), as well as gametophytic development in the moss *Physcomitrella patens* (Schuessele et al., 2016). Furthermore, protein degradation via this pathway plays a key role in the control of flooding tolerance (Gibbs et al., 2011; Licausi et al., 2011). Importantly, the Arg/N-degron pathway has also been implicated in plant defenses against pathogens (de Marchi et al., 2016; Gravot et al., 2016; Vicente et al., 2019), although a possible connection with AvrRpt2 activity has not been investigated.

In addition to RIN4, AvrRpt2 can cleave other host NOI domain-containing proteins with the AvrRpt2 consensus recognition motifs (Chisholm et al., 2005; Kim et al., 2005a; Takemoto and Jones, 2005; Afzal et al., 2011; Eschen-Lippold et al., 2016a). These NOI proteins have a similar domain architecture as RIN4 (Chisholm et al., 2005; Afzal et al., 2011) and are also presumed to be bound to the plasma membrane (Afzal et al., 2011; Afzal et al., 2013). Despite these similarities with RIN4, the role of the NOI domain proteins in either plant immunity or in promoting bacterial virulence following AvrRpt2 cleavage have remained largely elusive. In particular, although many of the NOI-domain protein fragments released upon AvrRpt2 cleavage are also predicted to bear destabilizing residues at their N-termini (Chisholm et al., 2005), several questions remain unanswered regarding the fate of these fragments: (i) are these fragments unstable in host cells following AvrRpt2 cleavage? (ii) is N-degron-mediated degradation required to regulate their abundance *in planta*?

Here, we show that native RIN4 fragments released after AvrRpt2 cleavage are unlikely to be N-degron substrates in a wild-type Arabidopsis background for RPS2. We also show, using selected NOI-domain proteins, that the C-terminal fragments released after AvrRpt2 cleavage, which start with a destabilizing residue, are typically unstable. We also reveal a role of the N-degron pathway in the degradation of NOI-domain proteolytic fragments generated after AvrRpt2 cleavage, so that several of these fragments are *bona fide* N-degron substrates. The latter results open new avenues of research to understand the role of AvrRpt2 in promoting bacterial virulence, as well as to dissect the role of the N-degron pathway in the regulation of the plant defense program in response to bacteria encoding the AvrRpt2 protease effector.

## Results

### A 12-mer RIN4-II N-terminal peptide is arginylated *in vitro*

Previously published experiments using a fusion protein composed of the first 30 amino acid residues of RIN4 fused to a C-terminal GFP reporter protein (RIN4^1-30^-GFP) suggested that cleavage of this RIN4 peptide at RCS1 resulted in the N-degron-mediated degradation of the resulting RIN4^11-30^-GFP fusion protein (Takemoto and Jones, 2005). The potential role of the N-degron pathway in clearing these fragments was also strengthened by the overall evolutionary conservation (with some exceptions; Supplemental Fig. S1) of the newly exposed destabilizing N-terminal residue in various RIN4 orthologs (Fig. 1C). N-degron-mediated degradation of the RIN4^11-30^-GFP fusion protein, and also of the native RIN4-II fragment, would require the deamidation of the newly exposed N-terminal Asn^11^ by the NTAN1 amidase into Asp, followed by arginylation by the Arg-transferases ATE1/2 (Fig. 1D). Before examining the potential degradation of the native RIN4-II and III fragments by the N-degron pathway *in planta*, we first tested if Arabidopsis ATE1 could arginylate the N-terminal sequence of RIN4-II using *in vitro* arginylation assays in conjunction with 12-mer peptides corresponding to amino acid residues 12 to 22 of RIN4 preceded by a variable N-terminal residue ‘X’ (X-^12^WEAEENVPYTA^22^). The residue ‘X’ was either an Asp residue (i.e. to mimic the suggested deamidation by NTAN1 of the newly exposed N-terminal Asn^11^ into Asp after AvrRpt2 cleavage), or a stabilizing residue Gly, which is not arginylated. As a control, we generated 12-mer peptides corresponding to the common N-terminal sequence of the RAP2.2 and RAP2.12 transcription factors, which are known Arg-transferase substrates following dioxygenation of their initial Cys residue (White et al., 2017). In these peptides, amino acid residues 3-13 of the RAP2.2/12 transcription factors was preceded by an N-terminal residue ‘X’ (X-^3^GGAIISDFIPP^13^), which is (i) an oxidized cysteine residue or sulfonic acid that can be arginylated; or (ii) unoxidized Cys (not recognized by Arg-transferases); or (iii) Asp; or (iv) Gly. As hypothesized, based on the known specificity of Arg-transferases in plants (Graciet et al., 2009; Graciet and Wellmer, 2010; White et al., 2017), the RAP2.2/12 peptides starting with oxidized Cys (sulfonic acid) or Asp were arginylated, in contrast to RAP2.2/12 peptides bearing unoxidized thiolated Cys or Gly (Fig. 1E). Importantly, in the same assays, the N-terminal RIN4-II peptide starting with an Asp residue was arginylated *in vitro*, whereas the same peptide bearing a Gly at the N-terminus was not (Fig. 1E). Our *in vitro* results hence suggest that the N-terminal region of the RIN4-II fragment can serve as an ATE1 substrate *in vitro*.

### AvrRpt2 cleavage of RIN4 *in planta* results in unstable fragments irrespective of the presence of a stabilizing or destabilizing N-terminal residue

In order to address the N-degron-mediated degradation of the native RIN4-II and RIN4-III fragments originating from AvrRpt2 cleavage *in planta*, we used RIN4-specific antibodies to track the fate of these fragments. We first tested if previously published RIN4-specific antibodies (Mackey et al., 2002; Liu et al., 2009) and a commercially available RIN4 antibody (aN-13, Santa Cruz Biotechnology Inc., Cat. No. sc-27369), were suitable to detect full-length RIN4, as well as the RIN4-II and III fragments (Fig. 2A). To this aim, we expressed in *E. coli* the RIN4-II and RIN4-III protein fragments, as well as full-length RIN4, as fusion proteins with a poly-histidine tag and compared the ability of the different RIN4-specific antibodies to recognize these fragments using immunoblotting (Supplemental Fig. S2). The results of these experiments indicated that previously published antibodies (designated as anti-RIN4#1 (Mackey et al., 2002) and anti-RIN4#2 (Liu et al., 2009)) were suitable to detect the full-length protein, as well as the RIN4-II and RIN4-III fragments, albeit with different sensitivities. The commercially available anti-RIN4#3 antibody (aN-13) detected both the full-length protein and the RIN4-II fragment, but not the C-terminal RIN4-III fragment, and hence was not suitable for our experiments. All subsequent immunoblots were carried out with the anti-RIN4#2 as the primary antibody, because it produced less background (data not shown).

**Figure 2:**
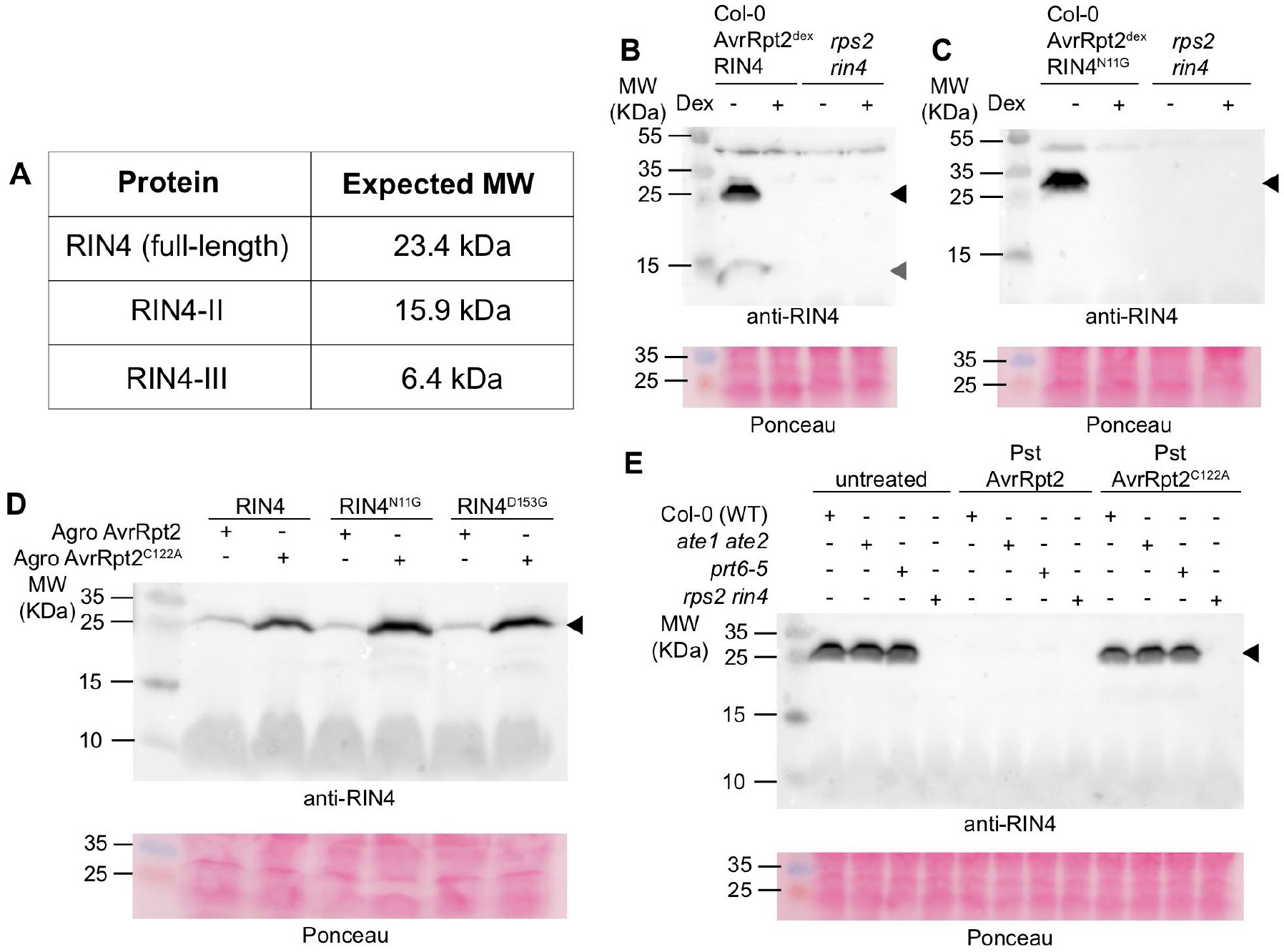
The instability of RIN4-II and RIN4-III proteolytic fragments does not depend on the N-degron pathway. **(A)** Predicted molecular weights (MW) of full-length RIN4 and proteolytic fragments released following AvrRpt2 cleavage. **(B)** Treatment with a dexamethasone-containing solution induces the disappearance of endogenous full-length RIN4 in Col-0 AvrRpt2^dex^ seedlings. Seedlings were grown in liquid 0.5xMS medium for 8 days and treated with a dexamethasone-containing solution for 5 hrs. Black arrowhead indicates full-length RIN4; grey arrowhead indicates a cross-reacting protein, which is also detected in *rin4 rps2* samples with longer exposure times (Supplemental Fig. S3). Data shown are representative of 3 independent replicates. **(C)** Induction of AvrRpt2 expression also triggers the disappearance of full-length RIN4^N11G^ in Col-0 AvrRpt2^dex^ 35S_pro_: RIN4^N11G^ seedlings. The Gly-RIN4-II fragment is not detected despite the presence of a stabilizing N-terminal residue. Seedlings were grown in liquid 0.5xMS medium for 8 days and treated with a dexamethasone-containing solution for 5 hrs. Black arrowhead indicates full-length RIN4. Data are representative of 3 independent replicates. **(D)** ^N11G^RIN4-II and ^D153G^RIN4-III fragments are not stabilized in tobacco. Four-week-old tobacco plants were co-infiltrated with *A. tumefaciens* strains coding for different versions of RIN4 and either an active or inactive variant of AvrRpt2. Tissue for immunoblot was collected 24 hrs after co-infiltration. Black arrowhead indicates full-length RIN4. Data is representative of 2 independent replicates. **(E)** RIN4-II and III fragments are not stabilized in mutant backgrounds for N-degron pathway enzymatic components. Plants of the indicated genotypes were left untreated or were inoculated with *Pst* AvrRpt2 or *Pst* AvrRpt2^C122A^ (5 × 10^7^ cfu/mL). Inoculated leaves were collected at 8 hpi for immunoblot analysis with the anti-RIN4#2 antibody. Black arrowhead indicates full-length RIN4. Data is representative of 3 independent replicates. For all panels, original data is presented in Supplemental Fig. S3.

To investigate the instability of the native RIN4 proteolytic fragments *in planta*, we conducted immunoblot experiments using protein extracts from wild-type seedlings that also encoded a dexamethasone-inducible version of AvrRpt2 (lines noted Col-0 AvrRpt2^dex^). In the absence of dexamethasone (i.e. mock treatment), the endogenous full-length RIN4 protein accumulated in the cells (Fig. 2A-B and Supplemental Fig. S3). In these conditions, a protein fragment of ~12 kDa was also detected by the anti-RIN4#2 antibody. However, longer exposure of the same immunoblot indicated that this protein was also detected in *rps2 rin4* double mutant plants (Supplemental Fig. S3), suggesting that it corresponds to a non-specific cross-reacting protein. When expression of AvrRpt2 was induced by dexamethasone treatment for 5 hrs, the full-length RIN4 protein was no longer detectable, as a result of its cleavage by AvrRpt2. In addition, the RIN4-II and RIN4-III fragments could not be detected, suggesting that the two proteolytic fragments were unstable (Fig. 2B and Supplemental Fig. S3). We next tested the importance of the newly exposed destabilizing residue (Asn^11^) of the RIN4-II fragment for degradation. To this aim, we generated stable Arabidopsis Col-0 AvrRpt2^dex^ lines that also expressed the full-length RIN4 in which Asn^11^ was substituted to Gly (RIN4^N11G^) under the control of the constitutive Cauliflower Mosaic Virus 35S promoter (lines designated as AvrRpt2^dex^ RIN4^N11G^). This mutation was unlikely to affect cleavage of RIN4 by AvrRpt2 because AvrRpt2 recognizes motifs with a Gly residue at the same location (Chisholm et al., 2005). While full-length RIN4^N11G^ accumulated in the absence of dexamethasone, the induction of AvrRpt2 expression led to a loss of full-length RIN4^N11G^, but not to the accumulation of the RIN4-II^N11G^ proteolytic fragment. The absence of RIN4-II^N11G^ detection suggests that the presence of an N-terminal stabilizing residue such as Gly failed to stabilize the RIN4-II fragment *in planta* (Fig. 2C).

To verify these results, we also conducted transient expression experiments in tobacco using co-infiltration with *A. tumefaciens* strains encoding AvrRpt2 or AvrRpt2^C122A^ (a catalytically inactive AvrRpt2 (Axtell et al., 2003)), and *A. tumefaciens* strains coding for different versions of the full-length RIN4 protein, including a wild-type version of the protein, a RIN4^N11G^ mutant or a RIN4^D153G^ mutant, which contains an Asp^153^ to Gly substitution at the N-terminus of the RIN4-III fragment generated by AvrRpt2 cleavage.

Immunoblot analysis using tissue collected 24 hrs after co-infiltration indicated that expression of neither RIN4^N11G^ nor RIN4^D153G^ impaired cleavage by AvrRpt2, as the full-length version of the protein decreased in abundance similarly to the wild type (Fig. 2D and Supplemental Fig. S3). Furthermore, accumulation of RIN4-II^N11G^ and RIN4-III^D153G^ fragments was not observed, suggesting that *in planta* destabilization of the full-length, untagged, RIN4-II and RIN4-III fragments released after AvrRpt2 cleavage does not require the presence of an N-terminal destabilizing residue.

### AvrRpt2 cleavage of RIN4 *in planta* results in unstable fragments in *Arabidopsis* N-degron pathway mutant backgrounds

To further test the potential N-degron-mediated degradation of both the RIN4-II and III fragments *in planta* with the least possible disruption to their conformation and expression levels, we assessed the stability of the native fragments released from endogenous RIN4 in wild-type and mutant backgrounds for the Arg-transferases (*ate1 ate2* mutant (Graciet et al., 2009; Holman et al., 2009)) and for the N-recognin PRT6 (*prt6-5* (Graciet et al., 2009)). If the RIN4-II and III fragments are indeed N-degron pathway substrates, we would predict them to be stabilized in the *ate1 ate2* and *prt6-5* mutants (Fig. 1D) after inoculation with a *Pseudomonas syringae* pv. tomato DC3000 (*Pst*) strain expressing AvrRpt2. As a control experiment for potential effects by other *Pst* (effector) proteins, we also inoculated the wild-type and mutant plants with a *Pst* strain containing the inactive AvrRpt2^C122A^ variant (Fig. 2E and Supplemental Fig. S3). Immunoblot analysis of the protein extracts indicate that inoculation with *Pst* AvrRpt2, but not with the *Pst* AvrRpt2^C122A^, results in disappearance of the full-length endogenous RIN4 protein as a result of AvrRpt2 activity at 8 hpi. However, neither the RIN4-II nor the RIN4-III fragments accumulated in the two mutant backgrounds.

Taken together, our data suggest that, *in planta*, destabilization of the RIN4-II and III fragments released after AvrRpt2 cleavage does not require the N-degron pathway.

### The relative lifetime of the RIN4-II and RIN4-III fragments is similar in *ate1 ate2* and wild-type plants

The experiments conducted above allowed us to monitor the accumulation of the RIN4-II and III fragments within a few hours after RIN4 cleavage by AvrRpt2. While a *bona fide* N-degron substrate should be stabilized in these conditions, the experimental approaches did not allow us to compare the degradation rate of the different fragments. To obtain a more accurate estimate of the relative lifetime of the RIN4-II and III fragments *in planta*, we used the tandem fluorescent timers technique (tFT) (Khmelinskii et al., 2012; Khmelinskii et al., 2016), which has recently been developed for plant-based systems ((Zhang et al., 2019); see companion manuscript). In this approach, a protein may be expressed as a fusion consisting of ubiquitin, followed by a residue ‘X’, the sequence of the protein/fragment of interest, mCherry and finally superfolder GFP (sfGFP). In these fusion proteins, the N-terminal ubiquitin moiety is cleaved off by deubiquitylating enzymes after the last residue of ubiquitin (Varshavsky, 2005), thus resulting in a X-protein-mCherry-sfGFP (X-protein-tFT) fusion protein with the residue X at the N-terminus. Measuring the ratio of the slow maturing mCherry to fast maturing sfGFP fluorescence intensities allows to study in a more dynamic manner the relative lifetime of the fusion protein. Importantly, this technique has been validated in Arabidopsis using the N-degron tFT reporter constructs Ub-M-tFT and Ub-R-tFT in the wild type and in a *prt6-5* mutant background (Zhang et al., 2019), showing that it is suitable to study N-degron-mediated degradation.

To apply the tFT approach, we generated constructs similar to those used in (Zhang et al., 2019), which allowed the expression of Ub-N-RIN4-II-tFT or Ub-D-RIN4-III-tFT fusions (Fig. 3A) under the control of the 35S promoter. After transient expression in leaves, the relative stability of the resulting N-RIN4-II-tFT and D-RIN4-III-tFT proteins was not significantly different in wild-type and *ate1 ate2* mutant backgrounds (Fig. 3B). These results hence further suggest that the RIN4-II and III fragments are not N-degron pathway substrates.

**Figure 3:**
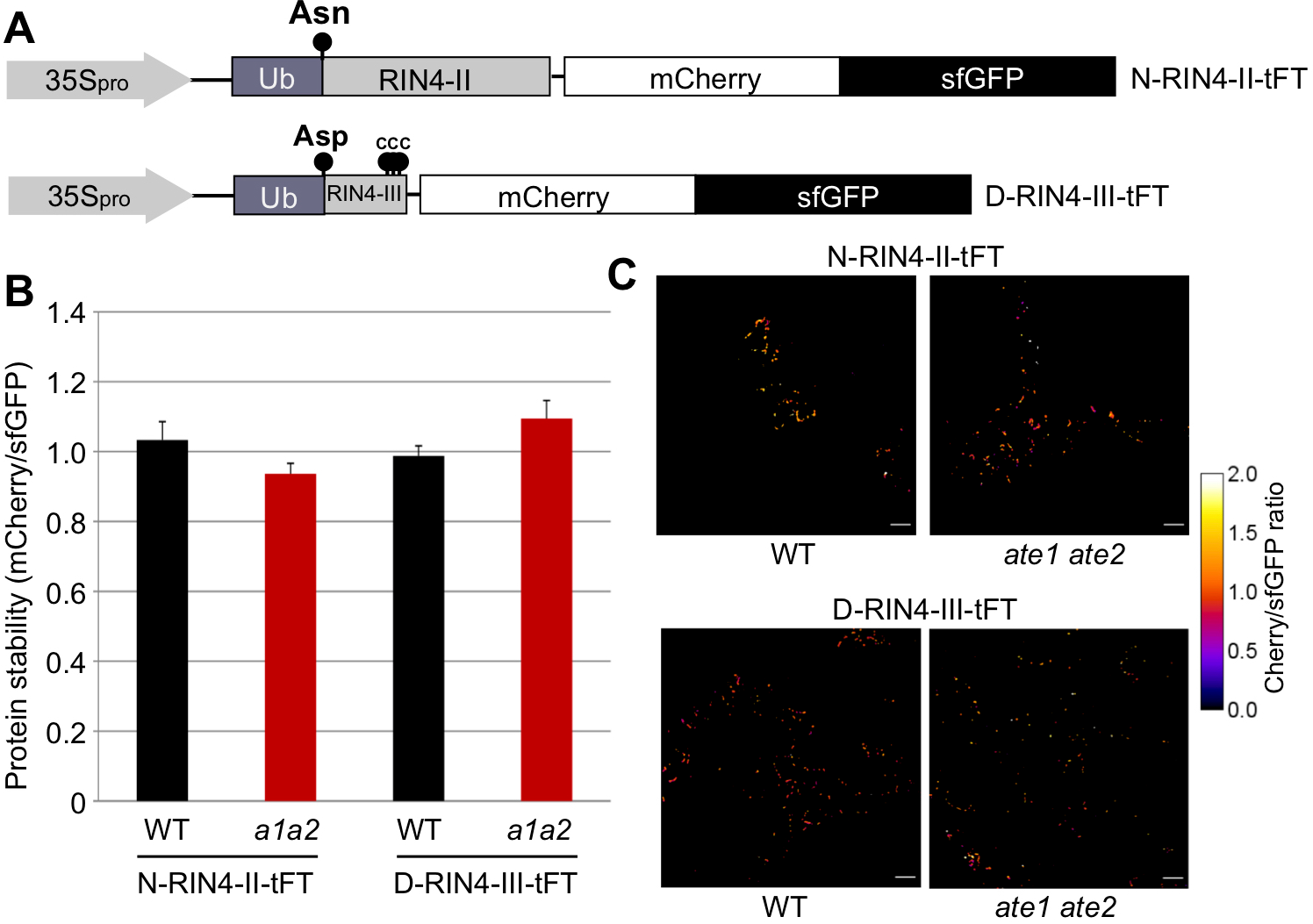
The relative lifetime of RIN4-II-tFT and RIN4-III-tFT fusion proteins is similar in wild-type and *ate1 ate2* plants. **(A)** Schematic representation of the ubiquitin fusion tFT constructs generated to determine the relative lifetime of the RIN4-II and III fragments. The N-terminal residue exposed after cleavage of ubiquitin are indicated (i.e. Asn for RIN4-II and Asp for RIN4-III). **(B)** Protein stability of the RIN4-II-tFT and RIN4-III-tFT fusion proteins in epidermal cells of 5-week old wild-type or *ate1 ate2* (*a1a2*) mutant plants, as determined by the ratio of mCherry (labelled Cherry) to sfGFP fluorescence intensity. Values represent mean and standard error of 5-7 independent replicates, and are not significantly different (One-way ANOVA). **(C)** Representative false-color images of Arabidopsis leaf epidermal cells expressing RIN4-II-tFT and RIN4-III-tFT reporters for calculation of mCherry/sfGFP ratios. The heatmap indicates the intensity ratio of mCherry to sfGFP, with blue corresponding to an unstable protein and white a stable one. Scale bar: 50 μm.

However, the N-RIN4-II-tFT and D-RIN4-III-tFT did not exhibit the expected cytosolic and plasma membrane localization (Fig. 3C and Supplemental Figs. S4 and S5), respectively (Kim et al., 2005a; Takemoto and Jones, 2005; Afzal et al., 2011), which may have impaired their potential N-degron-mediated degradation (see also Discussion).

### A subset of NOI C-terminal proteolytic fragments are unstable and accumulate upon mutation of the newly exposed N-terminal destabilizing residue

In addition to RIN4, AvrRpt2 has been shown to cleave other host NOI domain-containing proteins (Chisholm et al., 2005; Takemoto and Jones, 2005; Afzal et al., 2011; Eschen-Lippold et al., 2016a). AvrRpt2 cleavage of several NOI proteins is also predicted to generate proteolytic fragments starting with secondary destabilizing residues (Table 1), which could be sufficient to target them for degradation by the N-degron pathway, following arginylation and recognition by PRT6. To test the potential N-degron-dependent degradation of these C-terminal fragments, we selected for our analysis 6 different NOI proteins among the 14 NOI-domain proteins predicted to be encoded in the Arabidopsis genome. NOI1, NOI2, NOI3, NOI5, NOI6 and NOI11 were chosen to cover members of different clades (Eschen-Lippold et al., 2016a). NOI5, which is the closest in sequence to NOI1, was also included to test if closely related NOIs behave similarly. Next, we designed double-tagged versions of the 6 NOI proteins (Fig. 4A) to allow their expression as GFP-NOI-HA fusion proteins under the control of the constitutive 35S promoter.

**Table 1:** Several NOI proteolytic fragments generated after AvrRpt2 cleavage are potential N-degron substrates. Proteolytic fragments predicted to be released after AvrRpt2 cleavage of several NOI proteins (Chisholm et al., 2005) start with N-terminal destabilizing residues. All fragments would require the activity of the Arg-transferases and PRT6 for N-degron-mediated degradation. The corresponding gene numbers (AGI) and predicted newly exposed N-terminal residues after AvrRpt2 cleavage are indicated. *the representative protein model of NOI10 (At5g48657.1) is not cleaved by AvrRpt2 due to lack of the conserved cleavage site (Eschen-Lippold et al., 2016a).

| AGI | Protein Name | Predicted N-terminal residue<br>of AvrRpt2 cleavage product |
| --- | --- | --- |
| At3g25070 | RIN4-II fragment | Asn (N) |
| At3g25070 | RIN4-III fragment | Asp (D) |
| At5g64850 | NOI6 |  |
| At5g09960 | NOI7 |  |
| At5g48657 | NOI10* |  |
| At3g07195 | NOI11 |  |
| At5g63270 | NOI1 | Glu (E) |
| At5g40645 | NOI2 |  |
| At2g17660 | NOI3 |  |
| At5g55850 | NOI4 |  |
| At3g48450 | NOI5 |  |
| At5g18310 | NOI8 |  |

**Figure 4:**
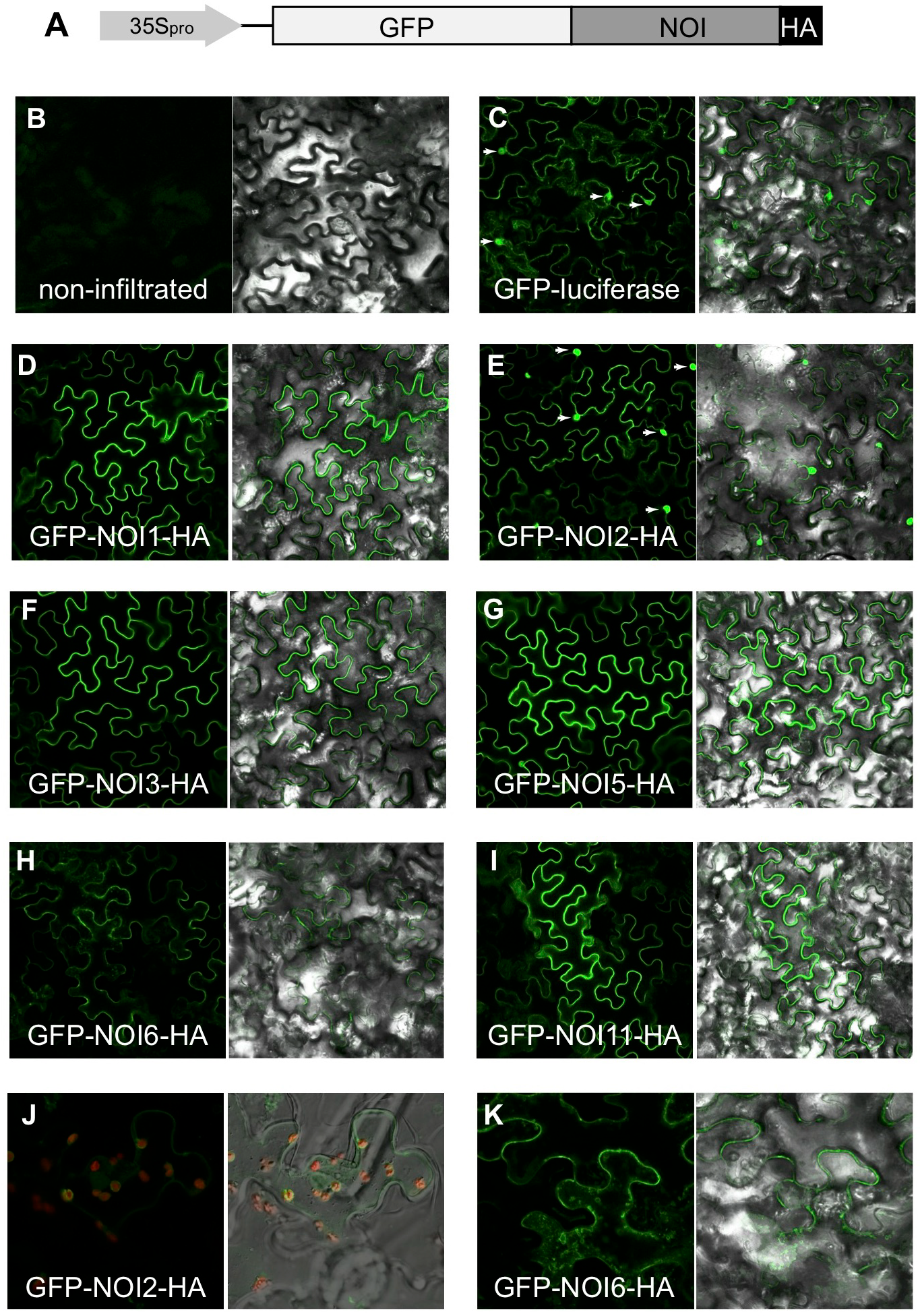
Subcellular localization of GFP-NOI-HA proteins. **(A)** Schematic representation of the double-tagged GFP-NOI-HA constructs under the control of the 35S promoter. **(B-I)** Confocal microscopy images of epidermal cells from tobacco leaves (abaxial side) 3 d after infiltration with *A. tumefaciens* coding for the different GFP-NOI-HA constructs. Putative nuclear signal is indicated with white arrowheads. **(J)** Confocal microscopy image of epidermal peel from tobacco leaf expressing GFP-NOI2-HA. Red signal corresponds to chlorophyll autofluorescence. **(K)** Confocal microscope image of epidermal cells from a tobacco leaf expressing GFP-NOI6-HA. Note the presence of intracellular vesicles in the presence of GFP-NOI6-HA. Green signal indicates GFP fluorescence.

We first determined the subcellular localization of the double tagged NOI proteins to check their predicted plasma membrane localization using transient expression in tobacco epidermal cells. A control GFP-luciferase fusion protein localized in the cytoplasm and in the nucleus of tobacco epidermal cells. In contrast, all GFP-NOI-HA proteins localized mostly to the periphery of epidermal cells (Fig. 4B-I), in agreement with their predicted subcellular localization (Afzal et al., 2013). The GFP-NOI2-HA fusion protein also displayed a strong nuclear signal. Epidermal peels were additionally used for closer inspection of the localization, where overlay of chlorophyll autofluorescence and GFP fluorescence revealed GFP-NOI2-HA signals surrounding the chloroplasts (Fig. 4J). In addition, the GFP-NOI6-HA fusion protein appeared to localize in small intracellular vesicle-like structures (Fig. 4K), which could be in agreement with its interaction with the exocyst complex subunits EXO70A1 and EXO70B1 (Afzal et al., 2013; Sabol et al., 2017). In sum, the GFP-NOI-HA-tagged proteins appear to localize as predicted based on their sequence and known protein interactors, suggesting that the double GFP/HA tags are unlikely to affect their sub-cellular localization.

We next tested if the GFP-NOI-HA fusion proteins were cleaved by AvrRpt2 and if the C-terminal ^Δ^NOI-HA fragments released after AvrRpt2 cleavage were unstable (Fig. 5A-C). To this aim, we infiltrated 4-week-old tobacco plants with different *A. tumefaciens* strains carrying a T-DNA coding for each of the GFP-NOI-HA constructs. After 72 hrs, the same leaves were infiltrated with either *Pst* AvrRpt2 or *Pst* AvrRpt2^C122A^. *Pst* infiltration was carried out 72 hrs after agroinfiltration to allow for sufficient accumulation of the GFP-NOI-HA fusion proteins, and tissue was collected 10 hrs after *Pst* inoculation. For each construct, immunoblots were performed with an anti-GFP and an anti-HA antibody in order to (i) verify the cleavage of the GFP-NOI-HA fusion proteins and (ii) test the stability of the N-terminal GFP-NOI^Δ^ and C-terminal ^Δ^NOI-HA fragments (Fig. 5C). Co-expression of the GFP-NOI-HA constructs and active AvrRpt2 resulted in the cleavage of all GFP-NOI-HA fusion proteins tested, as indicated by the disappearance of full-length fusion proteins concomitantly with the detection of an N-terminal GFP-NOI^Δ^ fragment in the presence of *Pst* AvrRpt2, but not *Pst* AvrRpt2^C122A^. Notably, the C-terminal ^Δ^NOI-HA fragments were not detected using an anti-HA antibody, demonstrating that these fragments are unstable in tobacco (Fig. 5C).

**Figure 5:**
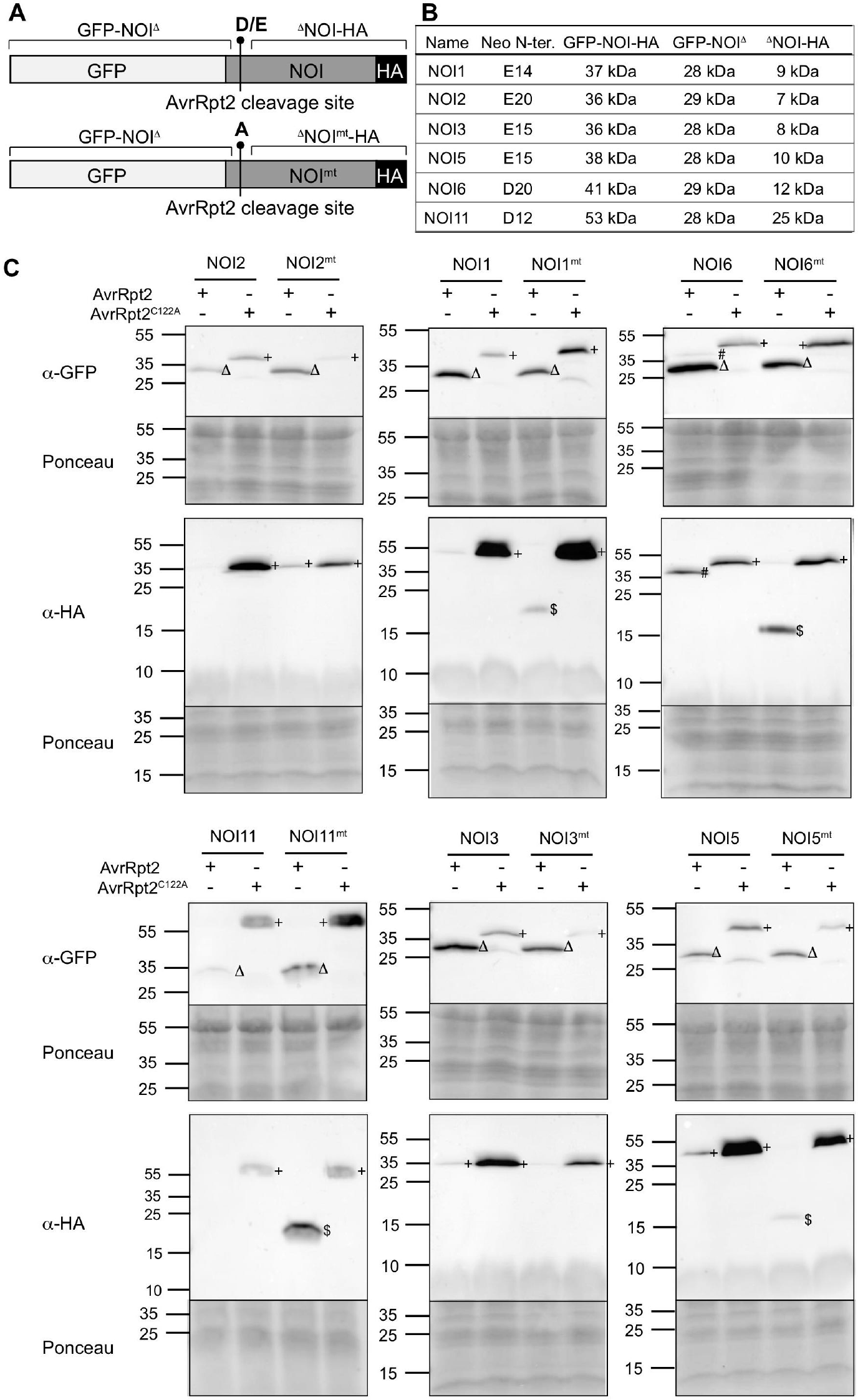
Stability of ΔNOI-HA and ΔNOI^mt^-HA fragments in tobacco. **(A)** Schematic representation of the GFP-NOI-HA and GFP-NOI^mt^-HA constructs. The predicted AvrRpt2 cleavage site is indicated, as well as the newly exposed Asp or Glu N-terminal residues (D/E; destabilizing residues) for wild-type sequences, or Ala (stabilizing residue) for the mutated version of the constructs (noted mt). GFP-NOI^Δ^refers to the N-terminally GFP-tagged fragments obtained after AvrRpt2 cleavage; ^Δ^NOI-HA and ^Δ^NOI^mt^-HA correspond to the C-terminal HA-tagged NOI fragments released after AvrRpt2 cleavage of the wild-type or mutated NOI proteins, respectively. **(B)** List of NOI proteins for which double-tag GFP-NOI-HA constructs were generated. The nature and position of the newly exposed N-terminal residue after AvrRpt2 cleavage is indicated (Neo N-ter.), as well as the calculated molecular weight of the different full-length fusion proteins and proteolytic fragments. **(C)** Stability of the GFP-NOI^Δ^, ^Δ^NOI-HA and ^Δ^NOI^mt^-HA fragments. Tobacco plants transiently expressing from the 35S promoter the GFP-NOI-HA or GFP-NOI^mt^-HA fusion proteins were inoculated with *Pst* AvrRpt2 or *Pst* AvrRpt2^C122A^. N-terminal fragments were detected using antibodies directed against the GFP tag, while C-terminal fragments were detected using anti-HA antibodies. +: full-length GFP-NOI-HA tagged proteins; Δ: GFP-NOI^Δ^fragments; $: ^Δ^NOI^mt^-HA fragments; #: aspecific cleavage product of GFP-NOI6-HA. These experiments were conducted three times independently with similar results. For all panels, original data is presented in Supplemental Fig. S6.

Because the ^Δ^NOI-HA fragments are predicted to start with a destabilizing residue, we tested a potential role of the N-degron pathway in their degradation by generating GFP-NOI^mt^-HA constructs, in which the newly exposed N-terminal destabilizing residue was changed into a stabilizing Ala residue (see Fig. 5B for specific mutations introduced in each of the NOI proteins). We then compared the accumulation of the ^Δ^NOI-HA and ^Δ^NOI^mt^-HA fragments. AvrRpt2 cleavage of GFP-NOI^mt^-HA fusion proteins was overall unaffected by the mutation introduced. Notably, the AvrRpt2-mediated cleavage of four GFP-NOI^mt^-HA proteins - specifically, NOI1, NOI5, NOI6 and NOI11 - generated detectable levels of the respective ^Δ^NOI^mt^-HA fragments in tobacco 10 hrs after *Pst* AvrRpt2 inoculation. This result suggests that NOI1, NOI5, NOI6 and NOI11, but not NOI2 and NOI3, could be potential N-end rule substrates.

To rule out any possible effects of other *Pst* DC3000 effector proteins on the processing and stability of the ^Δ^NOI-HA fragments, we conducted transient expression experiments in wild-type Arabidopsis protoplasts that were co-transfected with constructs coding for (i) wild-type AvrRpt2 or the inactive AvrRpt2^H208A^ variant (Cui et al., 2013); and (ii) the double-tagged GFP-NOI-HA or GFP-NOI^mt^-HA proteins. Proteins were then extracted for immunoblotting with anti-GFP and anti-HA antibodies. Similar to the results obtained in tobacco, co-expression of wild-type AvrRpt2 with either GFP-NOI-HA or GFP-NOI^mt^-HA resulted in cleavage of the fusion proteins (Fig. 6). Furthermore, the ^Δ^NOI-HA fragments released after cleavage of the wild-type double-tagged NOI proteins could not be detected in immunoblots with an anti-HA antibody, even after long exposure times (Fig. 6). In contrast, ^Δ^NOI^mt^-HA fragments accumulated to detectable levels with NOI1, NOI6 and NOI11, but not for NOI2, NOI3 and NOI5 (Fig. 6). Hence, irrespective of whether the AvrRpt2 effector was delivered via *Pst* or by direct AvrRpt2 expression in plant cells, our data suggest that the ^Δ^NOI^mt^-HA fragments obtained after cleavage of NOI1/6/11 could be N-degron pathway substrates. In contrast, fragments obtained after cleavage of NOI2/3/5 are not substrates in Arabidopsis.

**Figure 6:**
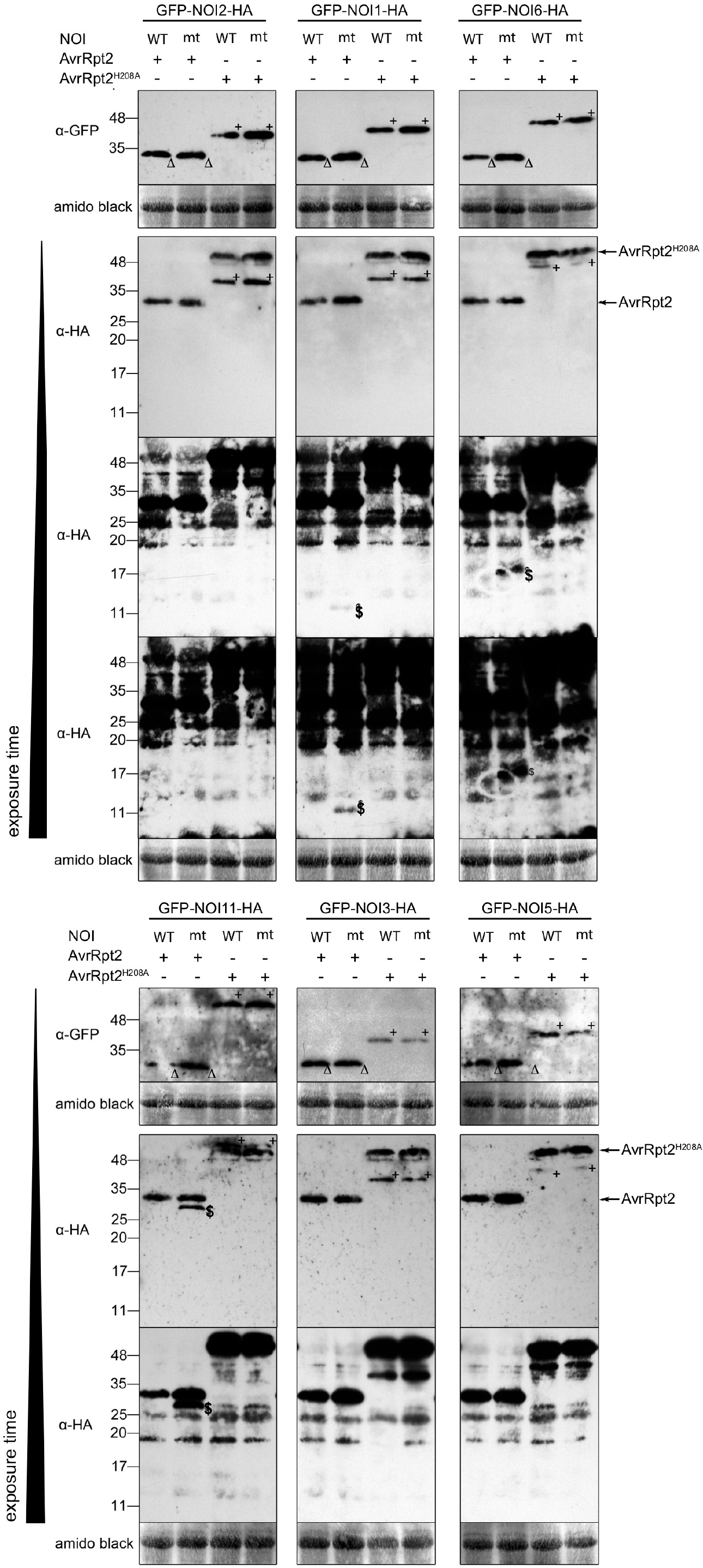
Stability of ^Δ^NOI-HA and ^Δ^NOI^mt^-HA fragments in wild-type Arabidopsis protoplasts. Stability of the GFP-NOI^Δ^, ^Δ^NOI-HA and ^Δ^NOI^mt^-HA fragments upon transient co-expression of GFP-NOI-HA and either wild-type AvrRpt2-HA or the AvrRpt2^H208A^-HA inactive variant. N-terminal fragments were detected using antibodies directed against the GFP tag, while C-terminal fragments were detected using anti-HA antibodies. For immunoblots with the anti-HA antibody, images of the same membrane with increasing exposure times are presented in order to show more clearly the stabilization of the different ^Δ^NOI^mt^-HA fragments. ‘+’: full-length GFP-NOI-HA tagged proteins; ‘Δ’: GFP-NOI^Δ^fragments; ‘$’: ^Δ^NOI^mt^-HA fragments; arrows indicate the self-processed AvrRpt2, as well as the unprocessed inactive AvrRpt2^H208A^ protease. For all panels, original data is presented in Supplemental Fig. S7. Data is representative of 3 independent replicates.

### Specific ^Δ^NOI-HA protein fragments constitute novel N-degron pathway substrates *in planta*

To further verify the N-degron-dependent degradation of the ^Δ^NOI-HA fragments, we conducted co-transfection experiments in protoplasts derived from wild-type (Col-0) plants, as well as from the *ate1 ate2* and *prt6-1* (Garzon et al., 2007) mutants, which are affected for Arg-transferases and PRT6 (Fig. 1A). The use of these mutant plants in conjunction with the GFP-NOI-HA constructs allowed us to rule out possible non-N-degron-dependent effects of the mutation introduced in the GFP-NOI^mt^-HA constructs on the stability of the resulting fragments after AvrRpt2 cleavage. Co-expression of either GFP-NOI2-HA or GFP-NOI5-HA with AvrRpt2 in *ate1 ate2* or *prt6-1* protoplasts confirmed that the ^Δ^NOI2-HA and ^Δ^NOI5-HA fragments were not N-degron pathway substrates. Interestingly, though, when the GFP-NOI3-HA fusion was co-expressed with AvrRpt2, the ^Δ^NOI3-HA fragment was stabilized in a *prt6-1* mutant background, suggesting that it may be an N-degron substrate in Arabidopsis (Supplemental Fig. S8). No stabilization was observed in *ate1 ate2*, likely due to the lower expression levels of the proteins in this mutant background. Finally, co-expression of the double-tagged NOI1, NOI6 and NOI11 proteins with either HA-tagged AvrRpt2 or AvrRpt2^H208A^ indicated that in the presence of AvrRpt2, but not AvrRpt2^H208A^, the ^Δ^NOI-HA fragments were unstable in the Col-0 background. In contrast, the ^Δ^NOI1-HA, ^Δ^NOI6-HA and ^Δ^NOI11-HA fragments were stabilized in the *ate1 ate2* and *prt6-1* mutant backgrounds (Fig. 7). This stabilization confirms that the ^Δ^NOI-HA fragments generated after cleavage of NOI1, NOI6 and NOI11 by AvrRpt2 are *bona fide* N-degron pathway substrates in Arabidopsis and hence constitute a novel set of proteins that are targeted for degradation *via* this ubiquitin-dependent pathway.

**Figure 7:**
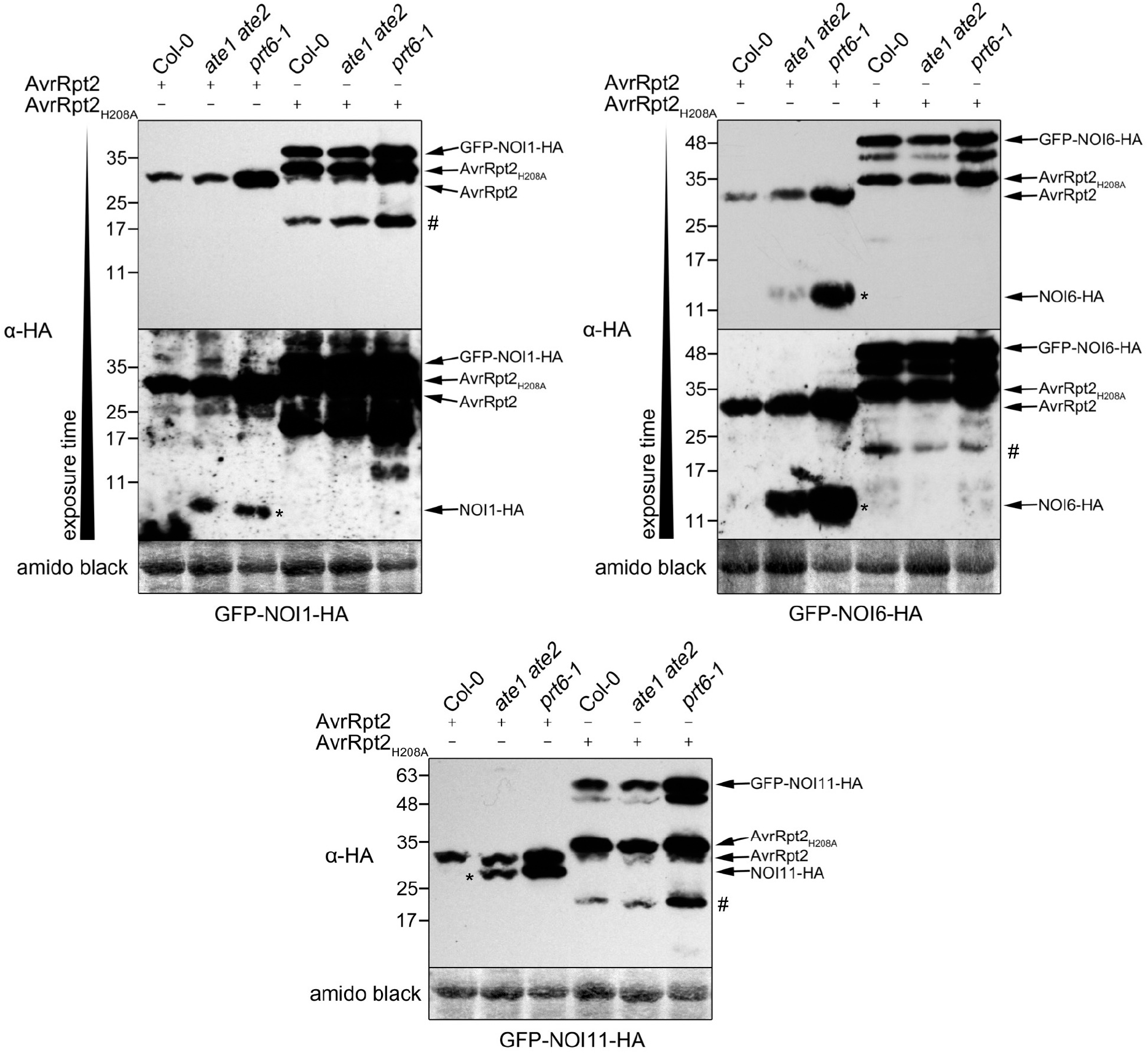
^Δ^NOI1-HA, ^Δ^NOI6-HA and ^Δ^NOI11-HA fragments are stabilized in *ate1 ate2* and *prt6-1* mutant protoplasts. To address the stability of wild-type ^Δ^NOI-HA fragments, protoplasts of wild-type Col-0 plants, as well as *ate1 ate2* and *prt6-1* mutant plants were co-transfected with plasmids coding for the GFP-NOI-HA fusion proteins (NOI1/6/11) and with plasmids encoding AvrRpt2-HA or AvrRpt2^H208A^-HA. ^Δ^NOI-HA C-terminal fragments (indicated with an asterisk) were detected using anti-HA antibodies. Images of the same membrane with increasing exposure times are presented in order to show more clearly the stabilization of the different ^Δ^NOI-HA fragments. ‘#’: aspecific band. For all panels, original data is presented in Supplemental Fig. S9. Data is representative of 3 independent replicates.

## Discussion

### Is the N-degron pathway involved in the degradation of the RIN4-II and RIN4-III fragments generated after AvrRpt2 proteolytic cleavage?

Cleavage of RIN4 by the effector protease AvrRpt2 and the resulting activation of RPS2 plays a key role in the onset of ETI. Several lines of evidence indicate that RIN4 fragments generated following AvrRpt2 cleavage are unstable (Axtell et al., 2003; Axtell and Staskawicz, 2003; Mackey et al., 2003), although several hours may be needed for their clearance (Kim et al., 2005a; Afzal et al., 2011). Importantly, the RIN4-II and III fragments that are released after AvrRpt2 cleavage bear Asn or Asp at their N-termini (Chisholm et al., 2005), respectively, both of which are destabilizing N-terminal residues (Graciet et al., 2010). Here, we used antibodies raised against RIN4 (Liu et al., 2009) to determine if the N-degron pathway could be responsible for the degradation of the native RIN4-II and RIN4-III fragments that are released upon cleavage by AvrRpt2. We combined different approaches, including (i) the expression of mutated versions of RIN4 in which the newly exposed N-terminal residues were changed to stabilizing ones; (ii) the monitoring of the RIN4-II and III fragments released after cleavage of the endogenous RIN4 protein in wild-type and N-degron mutant backgrounds; and (iii) the use of the recently developed tFT technique in plants (Zhang et al., 2019). The results of these experiments suggest that the RIN4-II and III fragments are degraded within a few hours of AvrRpt2 expression in Col-0 AvrRpt2^dex^ lines or in tobacco. However, contrary to what would be predicted if the fragments were N-degron substrates, the RIN4-II and III fragments did not accumulate to detectable levels in *ate1 ate2* mutant plants that lack Arg-transferase activity (Graciet et al., 2009).

Furthermore, mutation of the two newly exposed destabilizing N-termini (Asn^11^ and Asp^153^) into the stabilizing residue Gly was not sufficient to stabilize the RIN4-II and III fragments after cleavage of RIN4^N11G^ or RIN4^D153G^ by AvrRpt2 in wild-type Arabidopsis or tobacco plants. Altogether, our data suggest that a destabilizing N-terminal residue such as Asn or Asp is not necessary or sufficient for the degradation of the RIN4-II and III fragments, respectively. One possibility is that within the context of the native proteolytic fragments, the newly exposed N-terminal destabilizing residues may not be accessible for recognition by the Arg-transferases or by the cognate N-recognin PRT6. In the case of the RIN4-II fragment, which starts with N-terminal Asn, recognition by the amidase NTAN1 may also be problematic although spontaneous deamidation of Asn-initiated proteins to Asp was seen previously (Wang et al., 2009).

To compare more accurately the relative stability of the RIN4-II and III fragments, we generated tFT constructs (Zhang et al., 2019) that allowed the expression of RIN4-II-tFT and RIN4-III-tFT fusion proteins through the ubiquitin fusion technique (Varshavsky, 2005) instead of AvrRpt2 cleavage. The stability of the resulting tFT constructs (which are designed to bear N-terminal Asn or Asp, respectively) was examined in wild-type Col-0 and *ate1 ate2* mutant plants, and no differences in the relative stabilities of each of the fragments were found. The results of these experiments hence provide additional support for the idea that the N-degron pathway may not be required for the instability of the two fragments. Surprisingly though, the RIN4-II-tFT and RIN4-III-tFT did not exhibit the predicted cytosolic and plasma membrane localization, respectively (Kim et al., 2005a; Takemoto and Jones, 2005; Afzal et al., 2011). The tFT alone has a nucleo-cytoplasmic localization (Zhang et al., 2019), so that the punctuated localization of the RIN4-II-tFT and RIN4-III-tFT proteins is determined by the RIN4-II and III fragments. In the case of the RIN4-III-tFT fusion protein, it is possible that the presence of the C-terminal tFT tag may have affected the palmitoylation of the 3 C-terminal Cys residues of RIN4 (Fig. 1A), hence also preventing the plasma membrane tethering of RIN4-III-tFT. It is unclear though why the tFT fusion might have affected the cytosolic localization of the RIN4-II fragment, and whether this would have impaired the potential N-degron-mediated degradation of the fragments.

In sum, the data obtained *in planta* with full-length untagged RIN4 proteins strongly suggest that in plants in which RPS2 is functional, the native RIN4-II and RIN4-III fragments released after AvrRpt2 cleavage are not degraded by the N-degron pathway. Nevertheless, our data does not preclude a model whereby the RIN4-II and III fragments could be degraded through the recognition of a combination of N-terminal and internal degrons, as has been suggested for RAP2.12, which is targeted for degradation by the N-degron pathway, but has also been proposed to be a substrate of the E3 ubiquitin ligase SEVEN IN ABSENTIA of Arabidopsis thaliana2 (SINAT2) (Papdi et al., 2015). In the case of RIN4, the presence of an internal degron could still allow clearance of the RIN4 fragments in the absence of a destabilizing N-terminal residue.

### N-degron-mediated degradation of NOI proteins

Identification of the consensus sequence recognized and cleaved by AvrRpt2 in substrate proteins led to identification of novel putative AvrRpt2 substrates (Chisholm et al., 2005) belonging to the family of NOI proteins (Afzal et al., 2013). Subsequently, these NOI proteins were shown to be cleaved by AvrRpt2 (with the exception of NOI10) (Chisholm et al., 2005; Takemoto and Jones, 2005; Eschen-Lippold et al., 2016a). Notably, AvrRpt2 cleavage of several NOI proteins should result in protein fragments with a newly exposed destabilizing N-terminal residue (Table 1) (Chisholm et al., 2005; Takemoto and Jones, 2005). Based on their phylogenetic relationship (Eschen-Lippold et al., 2016a), we selected a set of 6 NOI proteins (NOI1/2/3/5/6/11) and tested (i) the stability of the C-terminal fragments released by AvrRpt2 cleavage; and (ii) the potential N-degron-mediated degradation of these fragments. Our experiments in tobacco and wild-type Arabidopsis protoplasts with double-tagged full-length NOI proteins (GFP-NOI-HA) indicate that the C-terminal fragments do not accumulate after cleavage by AvrRpt2. Introduction of point mutations to change the newly exposed N-terminal destabilizing residues into Ala, a stabilizing residue, was sufficient to stabilize the C-terminal fragments of NOI1/6/11. This would be consistent with their N-degron-mediated degradation.

To rule out the possibility that the single amino acid substitution introduced (Asp/Glu into Ala) could result in a conformational change that would prevent the degradation of the C-terminal fragments *via* an unknown internal degron, we also compared the stability of the wild-type HA-tagged NOI C-terminal fragments in wild-type and in mutant *ate1 ate2* protoplasts, following cleavage of the GFP-NOI-HA fusion proteins by AvrRpt2. The results of these experiments further confirmed that NOI1/6/11 constitute a novel set of N-degron substrates in plants. In addition, it is possible that the C-terminal fragment of NOI3 is also degraded by the N-degron pathway. The discovery of this novel set of N-degron substrates considerably expands the number of plant proteins that are targeted for degradation by the N-degron pathway. Thus far, the only known substrates of this pathway were a set of group VII Ethylene Response Factors (ERF-VII) transcription factors that act as master regulators of hypoxia response genes (Gibbs et al., 2011; Licausi et al., 2011) and VERNALIZATION2 (Gibbs et al., 2018), as well as BIG BROTHER (Dong et al., 2017).

The physiological relevance of the N-degron-mediated degradation of these NOI fragments now needs to be explored in the context of the different and varied functions of AvrRpt2, including its roles in virulence and in the onset of RPS2-mediated ETI. Considering the sequence similarities between the NOI domain of these proteins and the C-NOI domain of RIN4, it is possible that some of these fragments play a role in processes that are also regulated by RIN4, such as for example the repression of PTI. In this case, clearance of the C-terminal fragments by the N-degron pathway could provide a mechanism to alleviate the virulence-promoting activity of AvrRpt2 cleavage. Alternatively, cleavage of the NOI proteins could play a role in RIN4-independent functions of AvrRpt2, such as the repression of the flg22-induced activation of MPK4 and MPK11; the regulation of RPS2 signalling; or also in auxin-dependent mechanisms of AvrRpt2 induced virulence. However, the potential functional redundancy within the family of NOI proteins, which can be assumed based on the lack of phenotypes for single mutant plants (Eschen-Lippold et al., 2016a), makes it difficult to address this question using genetic approaches. Interestingly, not all NOIs may be cleaved by orthologs of AvrRpt2 encoded in the genomes of other bacteria, including some that are plant-associated rather than pathogenic (Eschen-Lippold et al., 2016a). It would hence also be interesting to study the role of the N-degron-mediated degradation of these NOI fragments in the context of the AvrRpt2 natural diversity.

### Determinants of N-degrons in plants

The results of our experiments with RIN4 and NOI proteolytic fragments highlight the difficulty of predicting and identifying N-degron substrates based on their sequence. Based on the known plant and mammalian N-degron substrates, it is thought that most N-degron pathway substrates are generated through proteolytic cleavage (Rao et al., 2001; Varshavsky, 2011; Piatkov et al., 2012; Dissmeyer et al., 2018). However, our results highlight the fact that knowledge of proteolytic cleavage sites and of the identity of the newly exposed N-terminal residue is insufficient to accurately predict N-degron-mediated degradation. There are classically 3 determinants associated with an N-degron (Bachmair and Varshavsky, 1989; Varshavsky, 2011): (i) a destabilizing residue at the N-terminus of the polypeptide chain; (ii) a Lys residue for ubiquitin conjugation; and (iii) sufficient flexibility of the polypeptide chain around the N-terminus or near the ubiquitylation site. The RIN4-II and III fragments, as well as the different NOI fragments generated after AvrRpt2 cleavage all have the required attributes of an N-degron substrate in that they bear a newly exposed N-terminal destabilizing residue after AvrRpt2 cleavage, several Lys residues are present in the C-terminal domains (between 8 and 12 Lys residues, except for RIN4-III, which only has 2 Lys residues), and all fragments are thought to be intrinsically unstructured (Sun et al., 2014; Lee et al., 2015a). Despite having these common attributes, only some of the fragments are N-degron pathway substrates. One possibility is that enzymatic components of the N-degron pathway may discriminate between these substrates based on the properties of the N-terminal region of the polypeptide chains. While the presence of an N-terminal destabilizing residue is crucial for N-degron-mediated degradation, the residue in second position is also known to influence the affinity or activity of both Arg-transferases and N-recognins towards their substrates (Choi et al., 2010; Matta-Camacho et al., 2010; Varshavsky, 2011; Wadas et al., 2016; Wang et al., 2018). While the exact effect of the amino acid residues neighboring the N-terminus on Arg-transferase and PRT6 specificity in plants are not known, the conservation of the first 2 residues (Asp or Glu, followed by Trp) in all fragments released after AvrRpt2 cleavage makes it unlikely to be the main determinant for selective degradation *via* the N-degron pathway. In addition, in the case of the RIN4-II fragment, whose 12-mer Asp-WEAEENVPYTA peptide that mimics the result of Asn^11^ deamidation by NTAN1 could be arginylated *in vitro*, it is also possible that the specificity of NTAN1 may preclude recognition of the newly exposed Asn^11^. However, very little is known about the influence of the sequence neighboring the N-terminal Asn for the specificity of NTAN1. Alternatively, the observed peptide arginylation may be the result of the *in vitro* approach, which further highlights the need to test substrate degradation *in planta*.

Notably, similar observations were made for the family of ERF-VII transcription factors. In particular, some of these ERF transcription factors in rice, namely ERF66, ERF67 and SUB1A-1, all have the attributes of an N-degron, as well as a conserved N-terminal region. Despite these common features, SUB1A-1 is not targeted for degradation by the N-degron pathway, while ERF66/67 are degraded *via* this pathway (Lin et al., 2019). The use of truncated proteins, as well as interaction assays, indicated that the C-terminal domain of SUB1A-1 could fold back onto the N-terminal region, thus precluding recognition of the N-terminal destabilizing residue by N-degron components (Lin et al., 2019). Interestingly, the N-terminal region of SUB1A-1 is also largely unstructured (Lin et al., 2019), similarly to the RIN4 fragments, and presumably also the NOI fragments. It is hence possible that differences in the conformation of the different fragments released after AvrRpt2 cleavage could explain the different degradation mechanisms. Alternatively, protein-protein interactions between the different NOI domains and other proteins may also prevent N-degron degradation depending on the interacting partner or the subcellular localization. The identification of protein interactors for both NOI-domain proteins and RIN4 already suggest protein-specific interactomes for each of these AvrRpt2 substrates (Liu et al., 2009; Afzal et al., 2013), which may correlate with differential recognition by N-degron components.

## Materials and Methods

### *In vitro* arginylation assays

The RIN4 and RAP2 peptides synthesized had the X-WEAEENVPYTA or X-GGAIISDFIPP sequence, respectively, and each peptide included a C-terminal PEG-biotin linker. For the arginylation assay, 1 µL of the purified His-ATE1 (corresponding to 0.5 µg of His-ATE1; cloning, expression and purification of His-ATE1 is described in Supplemental Materials) was incubated overnight at 30°C with 50 µL of the reaction mix (50 mM HEPES, pH 7.5; 25 mM KCl; 15 mM MgCl_2_; 100 mM DTT; 2.5 mM ATP; 0.6 mg/mL *E. coli* tRNA (Sigma); 0.04 µg/µL *E. coli* aminoacyl-tRNA synthetase (Sigma), 0.2 µCi ^14^C-Arg) and 50 µM of the indicated peptide substrates. The next day, for each reaction, 50 µL of avidin agarose bead slurry (Thermo Scientific) were equilibrated in PBS (100 mM NaH_2_PO_4_; 150 mM NaCl; 0.2% Nonidet P40; pH 7.2) and resuspended in 350 µL PBS. The reaction mixtures were added separately to 50 µL of avidin agarose bead and incubated at room temperature for 2 hrs with rotation. Subsequently, the beads were washed 4 times with 800 µL PBS. Finally, the beads were resuspended in 1 mL of a scintillation solution. Scintillation counting was performed in a total of 4 mL solution.

### Plant growth conditions

Tobacco plants were grown on a medium consisting of compost, perlite and vermiculite in a ratio of 5:2:3. The plants were grown under constant illumination at 20°C after being incubated at 4°C in the dark for 5 d.

### Treatment of plants with dexamethasone

Col-0 AvrRpt2^dex^ Arabidopsis seeds were grown in 3 mL liquid 0.5x MS medium (2.2 g/L MS salts, pH 5.7) for 7 or 8 d with shaking at ~130 rpm in continuous light. Dexamethasone (Sigma), prepared as a stock solution of 10 mM in ethanol, was added to the growth medium to a final concentration of 10 μM. Whole seedlings were collected 5 hrs after the addition of dexamethasone to the medium.

### *Pst* AvrRpt2 and *Pst* AvrRpt2^C122A^ inoculations of wild-type Col-0 and of *ate1 ate2* mutant plants for immunoblot analysis

*Pst* inoculations were performed on plants grown in Jiffy pots with a 9-hr light period and a light intensity of 190 µmol/(m^2^.s). *Pst* AvrRpt2 and *Pst* AvrRpt2^C122A^ were grown at 28°C on KB medium supplemented with 6 mM MgSO_4_. *Pst* strains were inoculated on 4-week-old plants following a high-humidity treatment 12 hrs before inoculation. A bacterial suspension at 5 × 10^7^ cfu/mL in 10 mM MgCl_2_ was infiltrated using a blunt syringe into the abaxial side of 3 leaves per plant. Three leaf discs (7 mm in diameter) from 3 different plants were collected at 8 hpi and pooled.

### Generation of RIN4 fragment tandem fluorescent timers and GFP-NOI-HA or GFP-NOI^mt^-HA-coding constructs

All cloning procedures are described in Supplemental Materials. Oligonucleotides used and plasmids generated are listed in Supplemental Tables 1 and 2, respectively.

### Tandem fluorescent timer experiments

Transient expression experiments with the RIN4-II-tFT and RIN4-III-tFT constructs were carried out using 5-week old Arabidopsis plants (Col-0 and *ate1 ate2*) using the protocol described in (Mangano et al., 2014). Confocal imaging was conducted 4 days after agroinfiltration. Arabidopsis leaves were covered with water and analyzed using the A1 confocal laser scanning microscope (Nikon). The fluorescence signal was imaged at 525/50 nm after excitation at 488 nm for sfGFP, and at 595/50 nm after excitation at 561 nm for mCherry. False color ratiometric images were generated after applying a Gaussian blur with a sigma of 1 and background subtraction in IMAGEJ (v.1.52h; https://imagej.nih.gov/ij). To quantify the mCherry to sfGFP ratio on a pixel by pixel basis, signal intensities of mCherry were divided by the intensities of the sfGFP using the image calculator of IMAGEJ. mCherry to sfGFP ratios were visualized by using the ImageJ lookup table ‘Fire’. Since the mCherry to sfGFP ratio is sensitive to any microscopic setting, only ratios generated with identical configuration were compared.

### Co-expression of RIN4 constructs with AvrRpt2 and AvrRpt2^C122A^

Co-expression of RIN4 constructs with AvrRpt2-FLAG or AvrRpt2^C122A^-FLAG was carried out in a similar manner as described (Day et al., 2005). Briefly, *Agrobacterium* strain C58 pGV2260 cells were transformed with the pMD-1 plasmids encoding the different RIN4 variants, AvrRpt2-FLAG or AvrRpt2^C122A^-FLAG and grown on LB agar with antibiotic selection for 3 d. After overnight growth in liquid culture with antibiotic selection, cells were collected by centrifugation and resuspended in infiltration medium (10 mM MES pH5.6, 10 mM MgCl_2_, 150 μM acetosyringone), to a final OD_600_ of 0.4 (RIN4 variants) and 0.1 (AvrRpt2-FLAG or AvrRpt2C122A-FLAG). Cells were syringe infiltrated into the abaxial side of tobacco leaves and tissue was collected 24 hrs post-infiltration.

### Transient gene expression in *N. benthamiana* followed by inoculation of *P. syringae*

*Agrobacterium* strain C58 pGV2260 cells were transformed with binary pMD-1 plasmids encoding RIN4 variants or with pML-BART plasmids encoding GFP-NOI-HA constructs and grown on LB agar with antibiotic selection for 3 d. After overnight growth in liquid culture with antibiotic selection, cells were collected by centrifugation and resuspended in infiltration media (10 mM MES pH5.6, 10 mM MgCl_2_, 150 μM acetosyringone), to a final OD_600_ of 0.75. Cells were syringe infiltrated into the abaxial side of tobacco leaves.

*Pseudomonas syringae* pathovar tomato DC3000 carrying a plasmid encoding AvrRpt2, or AvrRpt2^C122A^, or an empty plasmid were streaked onto an LB agar plate with 25 mg/L kanamycin, 5 mg/L tetracyclin, 100 mg/L rifampicin and grown for 2 d at 28°C. Cells from these plates were then resuspended in 10 mM MgCl_2_ to an OD_600_ of 0.7 before being infiltrated into previously agroinfiltrated *N. benthamiana* leaves using a 1 mL syringe. *Pst* infiltration was performed 72 hrs after the initial agroinfiltration. Tissue was collected and frozen in liquid nitrogen.

### Protein extraction and immunoblotting

To prepare total protein extracts from plant tissue (Arabidopsis or tobacco) for immunoblotting, plant tissue was ground to a fine powder in liquid nitrogen. This powder was then resuspended in 2x SDS loading buffer. Samples were spun at 18,000x g for 10 min and supernatant was transferred to a new 1.5 mL tube. This step was repeated before samples were placed at 95°C for 5 min. Samples were centrifuged at 18,000x g for 10 min and supernatant was used for subsequent analysis.

Primary antibodies used in this study for immunoblotting were anti-GFP (1:5,000; Abcam #Ab290), anti-HA (1:1,000; Sigma #H3663), anti-RIN4#1 (1:2,000; (Mackey et al., 2002)), antiRIN4#2 (1:2,000; (Liu et al., 2009)), anti-RIN4#3 (aN-13, 1:200; Santa Cruz # sc-27369). Secondary antibodies used were anti-rabbit HRP (A0545, Sigma), anti-mouse HRP (A9044, Sigma), both diluted 1:50,000. All antibodies were prepared in PBS-T supplemented with 5% milk powder (w/v).

### Plant growth conditions and transient expression in protoplasts

*Arabidopsis thaliana* wild type (Col-0), *prt6-1* and the double mutant *ate1 ate2* were grown on soil in a climate chamber for 5 to 6 weeks under controlled conditions (22°C, 8 hrs light / 16 hrs darkness; 140 µE). Isolation and transfection of protoplasts were performed as described in (Yoo et al., 2007). Protoplasts co-expressing AvrRpt2 (WT/H208A) and double-tagged NOI proteins (N-terminal GFP, C-terminal HA) were harvested by centrifugation after 16 hrs incubation at 18°C in the dark. Protoplast pellets were mixed with standard SDS-loading buffer, boiled for 5 min (95°C) and total protein samples were used for SDS-PAGE and immunoblotting. All antibodies were diluted in 1x TBS-T with 3% milk powder (primary antibodies: anti-GFP JL-8 (1:5,000), Takara Bio #632381; anti-HA.11 16B12 (1:1,000), Biozol #BLD-901515; secondary antibody: anti-mouse HRP (1:10,000), Sigma-Aldrich #A9044).

### Confocal imaging to visualize GFP-NOI-HA protein sub-cellular localization

*Agrobacterium* strain C58 pGV2260 cells were transformed with pML-BART vector encoding the different GFP-NOI-HA constructs and grown on LB agar with antibiotic selection for 3 d. After overnight growth in liquid culture with antibiotic selection, cells were collected by centrifugation and resuspended in infiltration medium (10 mM MES pH5.6, 10 mM MgCl_2_, 150 μM acetosyringone), to a final OD_600_ of 0.75. Cells were syringe infiltrated into the abaxial side of tobacco leaves. Three days after agroinfiltration an Olympus FluoView1000 laser scanning confocal microscope was used to visualize GFP fluorescence from the abaxial leaf side. A 488 nm excitation wavelength was used. GFP signal was collected at 500 - 550 nm and auto-fluorescence signal was collected at 600 - 700 nm. All imaging conditions were kept constant for all experiments.

## Funding information

KG, MS, RdM and this work were funded by grant 13/IA/1870 from Science Foundation Ireland to EG. LE-L is supported by grant #LE 2321/3-1 from the Deutsche Forschungsgemeinschaft (DFG) to JL. CN and MK were funded by a grant of the ScienceCampus Halle - Plant-Based Bioeconomy to ND and by grant #DI 1794/3-1 from the DFG. CN was also supported by a Ph.D. fellowship from the Landesgraduiertenförderung Sachsen-Anhalt. AK was supported by the ERASMUS+ exchange program. Research in MW group was funded by the (DFG) via the Collaborative Research Centre 1036 TP 13 and the European Union by the ERA-CAPS project KatNat.

## Author contributions

KG, AK and EG constructed plasmids used in tobacco and Arabidopsis protoplast experiments, including tFT plasmids; KG carried out tobacco transient expression experiments, immunoblots and confocal imaging, as well as all immunoblots for RIN4; LE-L conducted all NOI experiments in Arabidopsis protoplasts, including immunoblots; CN and MK constructed plasmids for *in vitro* arginylation assays and performed these assays; EL conducted tFT experiments; MS and RdM carried out inoculations with *Pst* AvrRpt2 and confocal analysis; EG, ND, JL and MW supervised the experiments; EG wrote the article with contributions of all the authors; EG agrees to serve as the author responsible for contact and ensures communication.

## Acknowledgements

We are very grateful to Stephen Chisholm and Brian Staskawicz for sharing the pMD-1 RIN4, pMD-1 RIN4^N11G^ and pMD-1 RIN4^D153G^ plasmids, the *Pst* AvrRpt2 and *Pst* AvrRpt2^C122A^ strains, as well as the *rps2 rin4*, Col-0 AvrRpt2^dex^ lines and the anti-RIN4#1 anti-serum. We are also very thankful to Gitta Coaker for sharing the purified RIN4-specific antibody (anti-RIN4#2), as well as the pMD-1 plasmids coding for AvrRpt2-FLAG or AvrRpt2^C122A^-FLAG. We wish to thank Dr. Ilona Dix (Maynooth University) for help provided with confocal imaging, Nicole Bauer (IPB Halle) for excellent technical assistance, and Brian Mooney (Maynooth University) for comments on the manuscript.

