## Supplemental Materials for "Differential N-end rule degradation of RIN4/NOI fragments generated by the AvrRpt2 effector protease"

##### Cloning of pDEST17 ATE1 to express His-tagged ATE1

The Arabidopsis ATE1 coding sequence was amplified using the primers ate1_tev_ss and ate1_as (see Supplemental Table 1 for oligonucleotide sequences). The ate1_tev_ss oligonucleotide included a sequence encoding the tobacco etch virus recognition sequence to create a cleavable linker to the N-terminal hexahistidine fusion for protein purification purposes. A subsequent PCR using the primers adapter and ate1_as was performed to provide attB-sites to use the PCR product in a BP reaction. The product was cloned into pDONR201 (Invitrogen) by BP cloning and into the destination vector pDEST17 (Invitrogen) by subsequent LR reaction, yielding pDEST17 ATE1.

##### Expression of His-tagged ATE1 in *E. coli*

The N-terminal hexahistidine-tagged ATE1 protein was expressed in BL21-CodonPlus (DE3)-RIL *E. coli* cells cultured in LB medium supplemented with 50 µg/mL carbenicillin. After cell growth to an $OD_{600}$ of 0.6, gene expression was induced with 1 mM isopropyl-β-D-thiogalactoside (IPTG), followed by culture for another 16 hrs at 18 °C. The cell pellet was resuspended in LEW buffer (50 mM $NaH_2PO_4$ pH8.0, 300 mM NaCl and 1 mM DTT). Lysis was performed by incubation with 1.2 mg/mL lysozyme for 30 min and subsequent sonification in the presence of 1 mM phenylmethylsulfonyl fluoride (PMSF). After purification by $Ni^{2+}$ affinity chromatography, the recombinant protein was subjected to Amicon Ultra-15 (30 K)

(Merck Millipore) filtration for buffer exchange to imidazole-free LEW containing 20% glycerol.

**Expression of His-tagged RIN4 and His-tagged RIN4 fragments in *E. coli***

*E. coli* BL21 Rosetta-GamiB (DE3) cells were transformed with pKG71, pKG72 or pKG73 and cells were grown overnight at 37°C on LB plates supplemented with 50 µg/mL kanamycin . The next day, individual colonies were used to inoculate liquid LB medium supplemented with 50 µg/mL kanamycin and grown at 37°C with shaking until $OD_{600}$ reached ~0.5. Cells were then chilled on ice for 30 min. To induce expression of recombinant proteins, isopropyl β-D-1-thiogalactopyranoside (IPTG) was added to a final concentration of 0.5 mM and cells were grown for 5 hrs at 30°C with shaking at 225 rpm. Cells were collected by centrifugation at 1,800x g for 10 min at 4°C and then stored at -80°C. To extract proteins from these pellets, cells were resuspended in 2x Laemmli buffer and placed at 95°C for 5 min. These samples were spun at 18,000x g for 10 min at room temperature and the supernatant was used for immunoblot analysis.

**Generation of RIN4 constructs for expression in *E. coli***

The cDNA sequence of RIN4 was PCR amplified with primers KG198/KG199 (full-length RIN4), KG20/KG201 (RIN4-II fragment) or KG202/KG203 (RIN4-III fragment) using Arabidopsis cDNA as a template. The resulting amplicons were digested with *Bam*HI and *Hin*dIII and inserted into pET28b plasmid that had been digested with the same enzymes. This yielded, respectively, plasmids pET28b RIN4 (pKG71), pET28b RIN4-II (pKG72) and pET28b RIN4-III (pKG73). See Supplemental Table 2 for a list of plasmids generated and used in this study.

**Generation of GFP-NOI-HA-coding constructs for expression in plants**

The NOI coding regions were amplified from Arabidopsis Col-0 cDNA using oligonucleotides indicated in Supplemental Table 1. The sequence coding for the HA tag was added during the PCR, using the ha6_lo oligonucleotide, which also included a *Sma*I restriction site. Each of the PCR products was digested with *Pst*I/*Sma*I and ligated into pBJ36 N6xOp cut with the same enzymes. This resulted in pBJ36 N6xOp:NOI-HA plasmids indicated in Supplemental Table 2.

To generate the pBJ36 35S:GFP-NOI-HA plasmids, the pBJ36 N6xOp:NOI-HA were digested *Pst*I, followed by end blunting with the T4 polymerase, and digestion with *Hin*dIII. The fragment of interest was then cloned into pBJ36 35S:GFP that had been digested with *Kpn*I, blunted with T4 polymerase and then digested with *Hin*dIII. Finally, the 35S:GFP-NOI-HA fragments were excised from the pBJ36 35S:GFP-NOI-HA plasmids (including 3'ocs terminator) by *Not*I digestion, and cloned into the binary vector pML-BART cut with *Not*I and dephosphorylated. This resulted in the pML-BART 35S:GFP-NOI-HA plasmids used for transient expression (Supplemental Table 2).

**Generation of GFP-NOI$^{mt}$-HA-coding constructs for expression in plants**

The pML-BART 35S:GFP-NOI$^{mt}$-HA were generated as indicated below:

- pMLBART 35S:GFP-NOI3$^{mt}$-HA (pKG61): pAK7 was used as a template for PCR reactions to mutagenize E15A using primers KG144/KG145 and KG146/KG147. The 2 PCR products were then fused by overlapping PCR using KG144/KG147. This product was digested using *Afl*II and *Xma*I and inserted into pBJ36 35S:GFP-NOI3-HA (pAK7) that had been digested with the same enzymes. The resulting construct (pKG56) was digested using *Not*I and the fragment of interest was inserted into pMLBART that had been digested with *Not*I and dephosphorylated.

- pMLBART 35S:GFP-NOI2$^{mt}$-HA (pKG74): pAK9 was used as a template for PCR reactions to introduce the E20A mutation using primers KG144/KG167 and KG168/KG147. The two PCR products were then fused by overlapping PCR using KG144/KG147. This product was digested using *Afl*II and *Xma*I and inserted into pAK9 that had been digested with the same enzymes. The resulting construct (pKG67) was digested with *Not*I and the fragment of interest inserted into pMLBART that had been digested with *Not*I and dephosphorylated.
- pMLBART 35S: GFP-NOI5$^{E15A}$-HA (pKG75): pAK8 was used as a template for PCR reactions to introduce the E15A mutation using primers KG144/KG169 and KG170/KG147. The two PCR fragments were then fused by overlapping PCR using KG144/KG147. This product was digested using *Afl*II and *Xma*I and inserted into pAK8 that had been digested with the same enzymes. The resulting construct (pKG68) was digested with *Not*I and the fragment of interest inserted into pMLBART that had been digested with *Not*I and dephosphorylated.
- pMLBART 35S: GFP-NOI6$^{mt}$-HA (pKG76): pAK6 was used as a template for PCR reactions to introduce the D20A mutation using primers KG144/KG171 and KG172/KG147. The two PCR fragments were then fused by overlapping PCR using KG144/KG147. This product was digested using *Afl*II and *Xma*I and inserted into pAK6 that had been digested with the same enzymes. The resulting construct (pKG69) was digested with *Not*I and the fragment of interest inserted into pMLBART that had been digested with *Not*I and dephosphorylated.

- pMLBART 35S: GFP-NOI11$^{mt}$-HA (pKG77): pAK19 was used as a template for PCR reactions to introduce the D12A mutation using primers KG144/KG173 and KG174/KG147. The two PCR fragments were then fused by overlapping PCR using KG144/KG147. This product was digested using *Afl*II and *Xma*I and inserted into pAK19 that had been digested with the same enzymes. The resulting construct (pKG70) was digested with *Not*I and the fragment of interest inserted into pMLBART that had been digested with *Not*I and dephosphorylated.
- pMLBART 35S: GFP-NOI1$^{mt}$-HA (pKG78): pAK5 was used as a template for PCR reactions to to introduce the E14A mutation using primers KG144/KG165 and KG166/KG147 that were then fused by overlapping PCR using KG144/KG147. This product was digested using *Afl*II and *Xma*I and inserted into pAK5 that had been digested with the same enzymes. The resulting construct (pKG66) was digested with *Not*I and the fragment of interest inserted into pMLBART that had been digested with *Not*I and dephosphorylated.

**Generation of RIN4 fragment tandem fluorescent timers**

The wild type ubiquitin-coding sequence was amplified using pKG30, a derivative of pEG368 (pML-BART Ub-R-LUC; (Graciet et al., 2010)), as a template for PCR with primers KG185/KG186 (RIN4-II) or KG185/KG191 (RIN4-III). RIN4 fragments were PCR amplified using Arabidopsis cDNA with primers KG187/KG188 (RIN4-II) or KG192/KG193 (RIN4-III). PCR products were fused using overlapping PCR with primers KG185 and KG188 (RIN4-II) or KG185/KG193 (RIN4-III) and cloned into a pJET plasmid using the pJET cloning kit (Thermo Fisher). The resulting pJET plasmids were then used as a template for a PCR reaction using primers KG185/KG220 (RIN4-II) or KG185/KG219 (RIN4-III), respectively. The resulting amplicon was digested with *Sma*I and cloned into pBin 35S:mCherry-GFP (Zhang et al., 2019) that had been digested with *Kpn*I and *Xho*I before being blunt-ended with T4 polymerase and dephosphorylated.

#### Supplemental Table 1: list of oligonucleotides used

| Oligo# | Sequence (5' -> 3') |
|---|---|
| **ate1_tev_ss** | GCTTAGAGAATCTTTATTTTCAGGGGATGTCTTTGAAAAACGATGCGAGT |
| **ate1_as** | GGGGACCACTTTGTACAAGAAAGCTGGGTATCAGTTGATTTCATACACCATTCTC TC |
| **primer adapter** | GGGGACAAGTTTGTACAAAAAAGCAGGCTTAGAGAATCTTTATTTTCAGGGG |
| **At44** | GAACACTGCAGATGGCGGAAAACAAAGGGAA |
| **At45** | GAACGTCGTATGGGTAGCCTGAACCACGGAAGCAAAACCATTTGT |
| **At42** | GAACACTGCAGATGGCATCGAATAATCAACAACG |
| **At43** | AACGTCGTATGGGTAGCCTGAACCGAAACAGCAGAACCGTTTCTT |
| **At36** | GAACACTGCAGATGGCGTCGAATGAAGCTGG |
| **At37** | GAACGTCGTATGGGTAGCCTGAACCAGCAAAAGTGAAACAGAGCC |
| **At38** | GAACACTGCAGATGGCAACGGGAAATAGAGCG |
| **At39** | AACGTCGTATGGGTAGCCTGAACCACGAAAACAAAGCCATTTCTT |
| **At46** | GAACACTGCAGATGGAAGATCGAAAAGAGAACAAGA |
| **At47** | AACGTCGTATGGGTAGCCTGAACCAGCTTTAACGCAACAGTTGAA |
| **At34** | CCACGCTGCAGATGGCAAACCGTCCACACGTTCC |
| **At35** | GAACGTCGTATGGGTAGCCTGAACCATATTTACTTCCCCTGCGGC |
| **ha6_lo** | TCGAACCCGGGTCACGCATAGTCAGGAACGTCGTATGGGTAGCCT |
| **KG144** | CGAGCTTAAGGGAATCGATTTCAA G |
| **KG145** | CATTCACATCCCATGCCCCGAAT |
| **KG146** | ATTCGGGGCATGGGATGTGAATG |
| **KG147** | CGAACCCGGGTCACGCAT |
| **KG167** | TCACGTCCCATGCTCCAAATTT |
| **KG168** | AAATTTGGAGCATGGGACGTGA |
| **KG169** | GCATCCCATGCCCCAAATTTTG |
| **KG170** | CAAAATTTGGGGCATGGGATGC |
| **KG171** | TTTTTGATCCCAAGCTCCAAACTGT |
| **KG172** | ACAGTTTGGAGCTTGGGATCAAAAA |
| **KG173** | TTGTTCCAGGCCCCAAACTTC |
| **KG174** | GAAGTTTGGGGCCTGGAACAA |
| **KG165** | ACATCCCATGCCCCAAACTT |
| **KG166** | AAGTTTGGGGCATGGGATGT |
| **KG198** | AAAAGGATCCGATGGCACGTTCGAATGTACC |
| **KG199** | AAAAAAGCTTTCATTTTCCTCCAAAGCCAAAG |
| **KG200** | AAAAGGATCCGAACTGGGAAGCTGAGGAGA |
| **KG201** | AAAAAAGCTTTCAACCGAATTTAGGCACCACT |
| **KG202** | AAAAGGATCCGGACTGGGACGAGAACAACC |
| **KG203** | AAAAAAGCTTTCATTTTCCTCCAAAGCCAAAG |
| **KG185** | AAAACCCGGGATGGAAATCTTCG |
| **KG186** | TCTCCTCAGCTTCCCAGTTACCACCTCTTAACCTGAGAAC |
| **KG191** | GGTTGTTCTCGTCCCAGTCACCACCTCTTAACCTGAGAAC |
| **KG187** | AACTGGGAAGCTGAGGAGAA |
| **KG188** | AAAACCCGGGGAACCGAATTTAGGCACCACTG |
| **KG192** | GACTGGGACGAGAACAACC |
| **KG193** | AAAACCCGGGGATTTTCCTCCAAAGCCAAAGC |
| **KG220** | AAAACCCGGGGACACCGAATTTAGGCACCACTG |
| **KG219** | AAAACCCGGGTGATTTTCCTCCAAAGCCAAAGC |

**Supplemental Table 2: List of plasmids generated for this study**

| Plasmid name | Plasmid number | Reference |
|---|---|---|
| pET28b RIN4 | pKG71 | This study |
| pET28b RIN4-II | pKG72 | This study |
| pET28b RIN4-III | pKG73 | This study |
| pBJ36 N6xOp:NOI1-HA | pEG273 | This study |
| pBJ36 N6xOp:NOI2-HA | pEG271 | This study |
| pBJ36 N6xOp:NOI3-HA | pEG276 | This study |
| pBJ36 N6xOp:NOI5-HA | pEG270 | This study |
| pBJ36 N6xOp:NOI6-HA | pEG274 | This study |
| pBJ36 N6xOp:NOI11-HA | pEG275 | This study |
| pBJ36 35S:GFP-NOI1-HA | pAK5 | This study |
| pBJ36 35S:GFP-NOI2-HA | pAK9 | This study |
| pBJ36 35S:GFP-NOI3-HA | pAK7 | This study |
| pBJ36 35S:GFP-NOI5-HA | pAK8 | This study |
| pBJ36 35S:GFP-NOI6-HA | pAK6 | This study |
| pBJ36 35S:GFP-NOI11-HA | pAK4 | This study |
| pML-BART 35S:GFP-NOI1-HA | pAK23 | This study |
| pML-BART 35S:GFP-NOI2-HA | pAK22 | This study |
| pML-BART 35S:GFP-NOI3-HA | pAK20 | This study |
| pML-BART 35S:GFP-NOI5-HA | pAK21 | This study |
| pML-BART 35S:GFP-NOI6-HA | pAK24 | This study |
| pML-BART 35S:GFP-NOI11-HA | pAK19 | This study |
| pBJ36 35S:GFP-NOI1$^{mt}$-HA | pKG66 | This study |
| pBJ36 35S:GFP-NOI2$^{mt}$ -HA | pKG67 | This study |
| pBJ36 35S:GFP-NOI3$^{mt}$ -HA | pKG56 | This study |
| pBJ36 35S:GFP-NOI5$^{mt}$ -HA | pKG68 | This study |
| pBJ36 35S:GFP-NOI6$^{mt}$ -HA | pKG69 | This study |
| pBJ36 35S:GFP-NOI11$^{mt}$ -HA | pKG70 | This study |
| pML-BART 35S:GFP-NOI1$^{mt}$-HA | pKG78 | This study |
| pML-BART 35S:GFP-NOI2$^{mt}$ -HA | pKG74 | This study |
| pML-BART 35S:GFP-NOI3$^{mt}$ -HA | pKG61 | This study |
| pML-BART 35S:GFP-NOI5$^{mt}$ -HA | pKG75 | This study |
| pML-BART 35S:GFP-NOI6$^{mt}$ -HA | pKG76 | This study |
| pML-BART 35S:GFP-NOI11$^{mt}$ -HA | pKG77 | This study |
| pBIN35S:Ub-RIN4-II-tFT | pKG79 | This study |
| pBIN35S:Ub-RIN4-III-tFT | pKG81 | This study |
| pDEST17 ATE1 | | This study |
| pUGW14-avrRpt2-HA | | (Eschen-Lippold et al., 2016) |
| pHBT-avrRpt2$^{H208A}$-HA | | (Cui et al., 2013) |

### Supplemental Figures

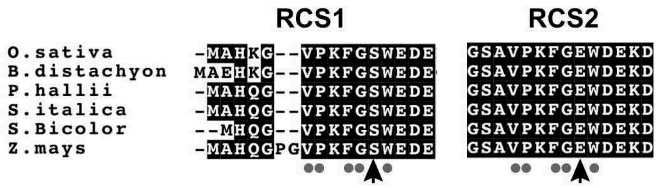


**Figure S1: Alignment of AvrRpt2 cleavage sites in RIN4 from monocots.** In monocots, some RIN4 orthologs bear Ser, a stabilizing residue, at the N-terminus of the RIN4-II fragment (black arrow) that is released after cleavage at RCS1. The newly exposed destabilizing residue (black arrow) after cleavage at RCS2 is conserved. Monocot RIN4 orthologs were identified using BLASTp with the sequence from *Z. mays* as a query. *O. sativa*: XP_015628179.1; *S. bicolor*: XP_021321828.1 (isoform X2); *B. distachyon*: XP_010233905.1 (isoform X2); *P. hallii*: XP_025793936.1; *S. italica*: XP_004981050.2 (isoform X2); *Z. mays*: AQK61593.1.

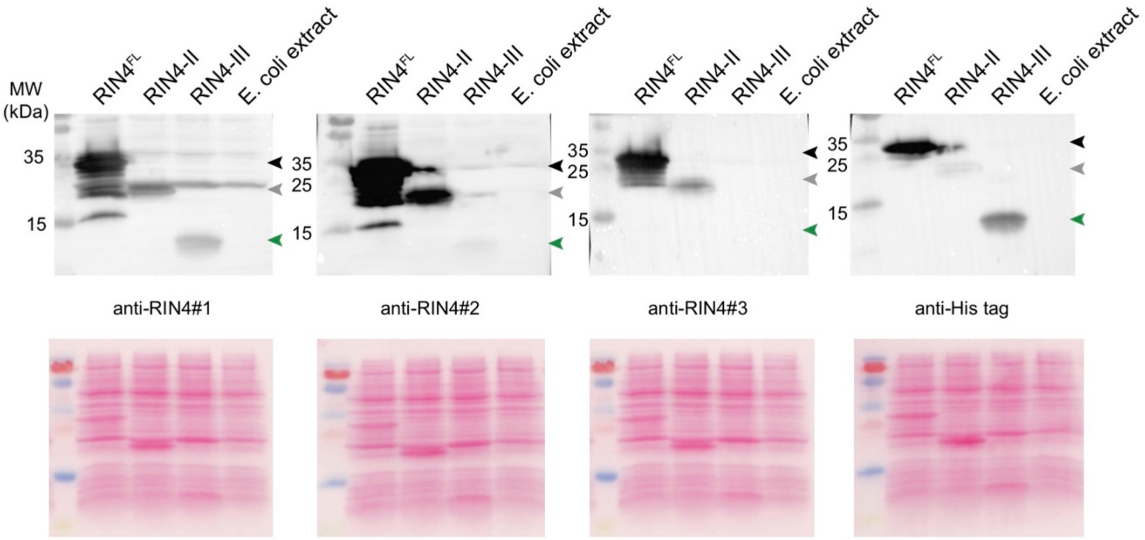


**Figure S2: Specificity of different antibodies towards RIN4 fragments and the full-length protein.** Crude lysates of *E. coli* BL21 Rosetta-GamiB (DE3) expressing 6xHis-RIN4 (~27 kDa; black arrowhead), 6xHis-RIN4-II (~19.5 kDa; grey arrowhead), or 6xHis-RIN4-III (~9.9 kDa; green arrowhead) were prepared in 2x SDS loading buffer. Protein concentration was measured using an amido black assay. Proteins were then separated using 14% SDS-PAGE electrophoresis and then analyzed using immunoblotting with the different RIN4-specific antibodies. An *E. coli* lysate generated from untransformed BL21 Rosetta-GamiB (DE3) was also analyzed to determine possible *E. coli* cross-reacting proteins for each of the antibodies tested. The RIN4-specific antibodies used correspond to anti-RIN4#1 (Mackey et al., 2002) and anti-RIN4#2 (Liu et al., 2009) and anti-RIN4#3 (aN-13; Santa Cruz).

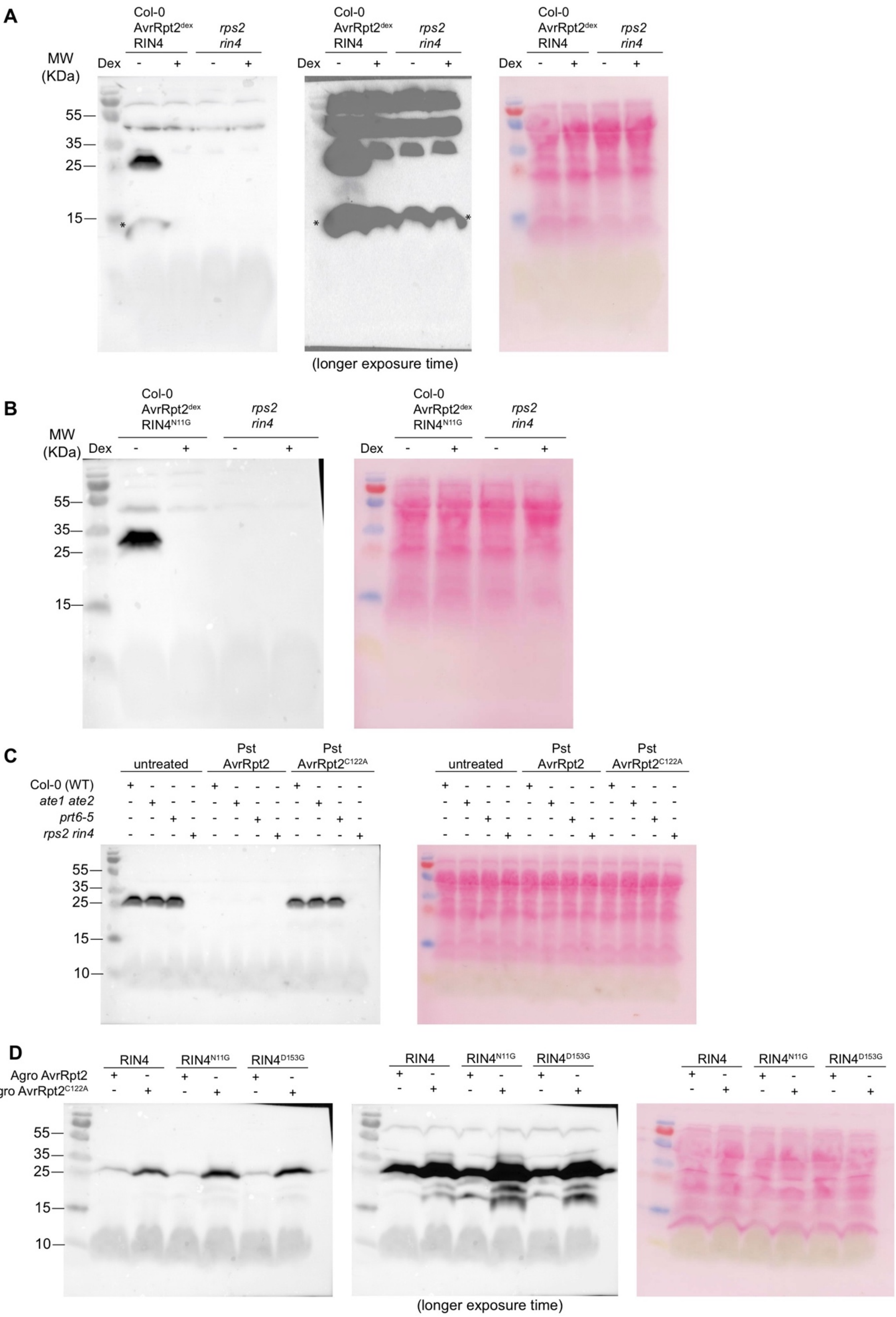


**Figure S3: Uncropped immunoblots shown in Fig. 3.** Ponceau-stained membranes and immunoblots are shown. In panels (A) and (D) longer exposure times are also shown.

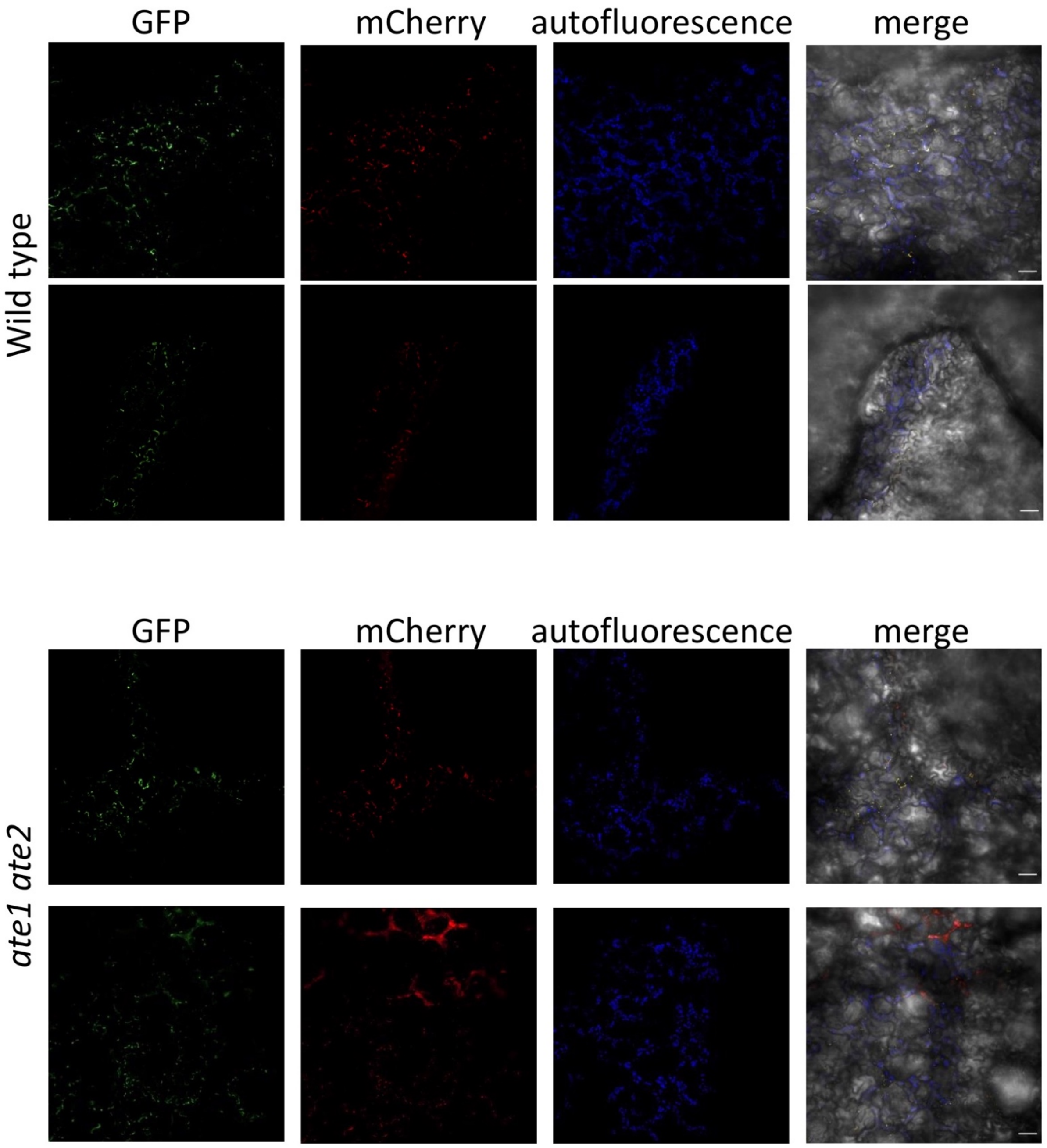


**Figure S4: Representative false-color images of cells transiently expressing N-RIN4-II-tFT in wild-type or in *ate1 ate2* plants.** *Agrobacterium*-mediated transient expression was carried out in 5-week-old plants as indicated in Materials and Methods. Scale bar 50 µm.

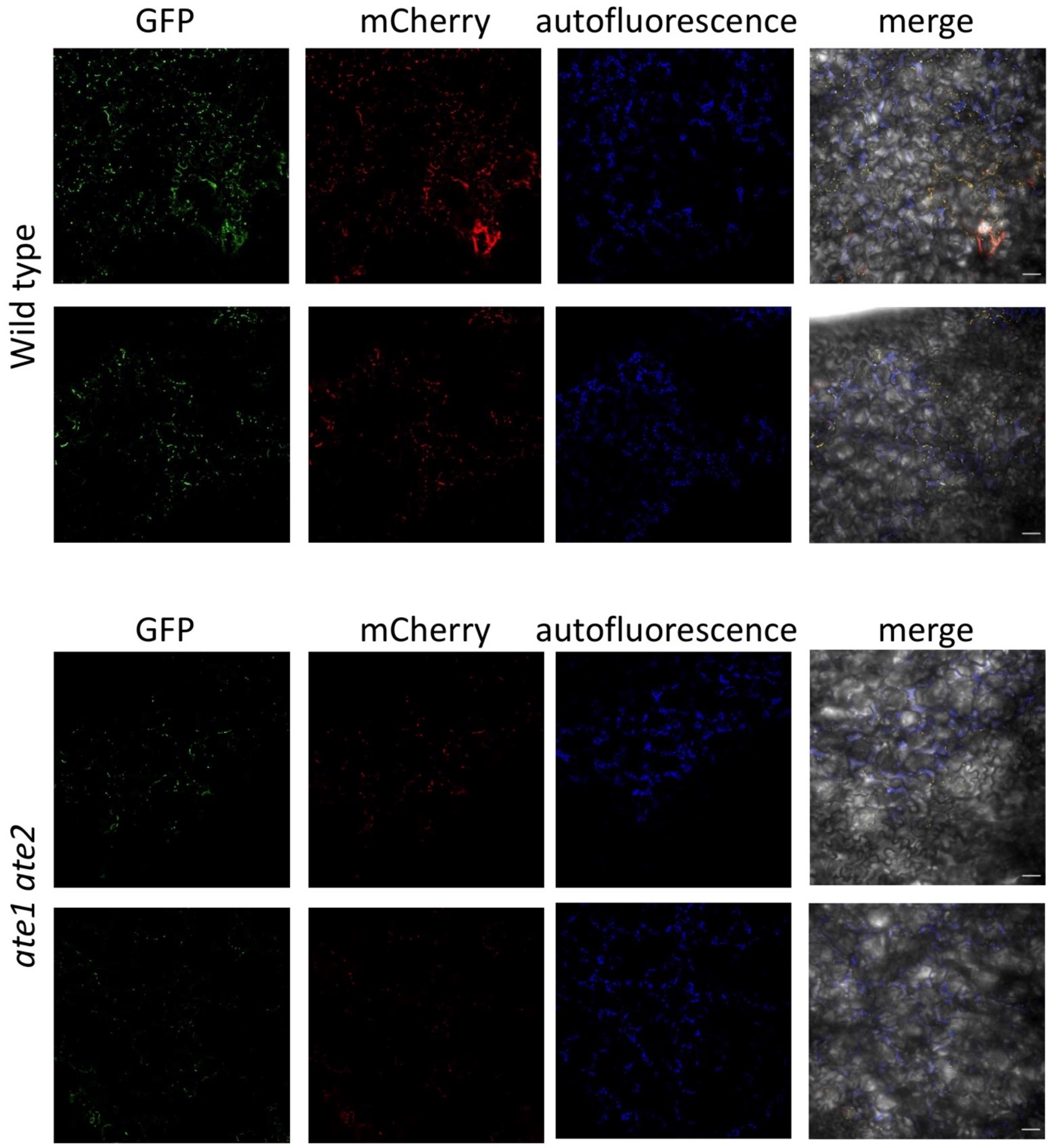


**Figure S5: Representative false-color images of cells transiently expressing D-RIN4-III-tFT in wild-type or in *ate1 ate2* plants.** *Agrobacterium*-mediated transient expression was carried out in 5-week-old plants as indicated in Materials and Methods. Scale bar 50 μm.

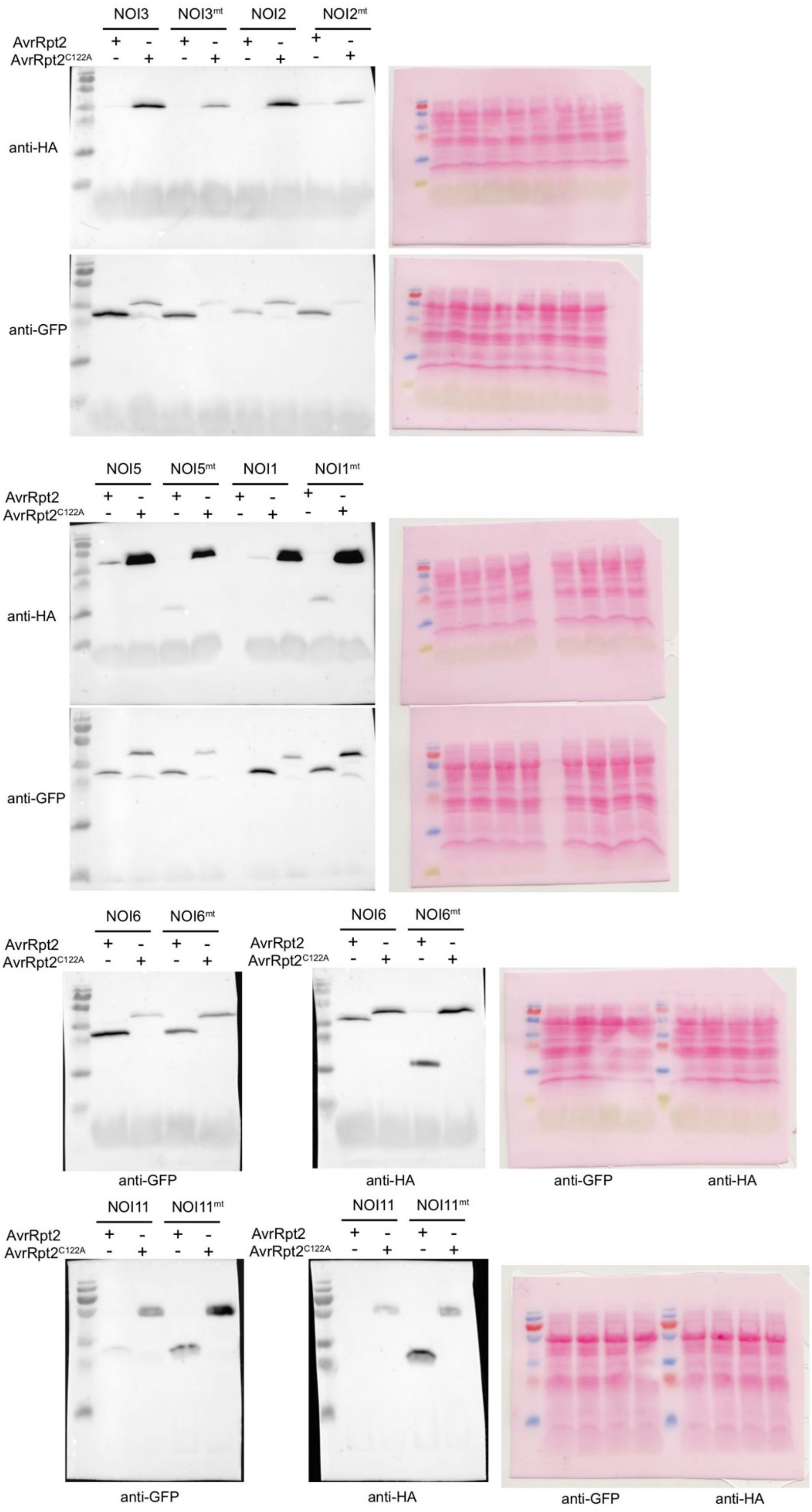


**Figure S6: Uncropped immunoblots shown in Fig. 5.** Ponceau-stained membranes and immunoblots are shown. Data is representative of 3 independent replicates.

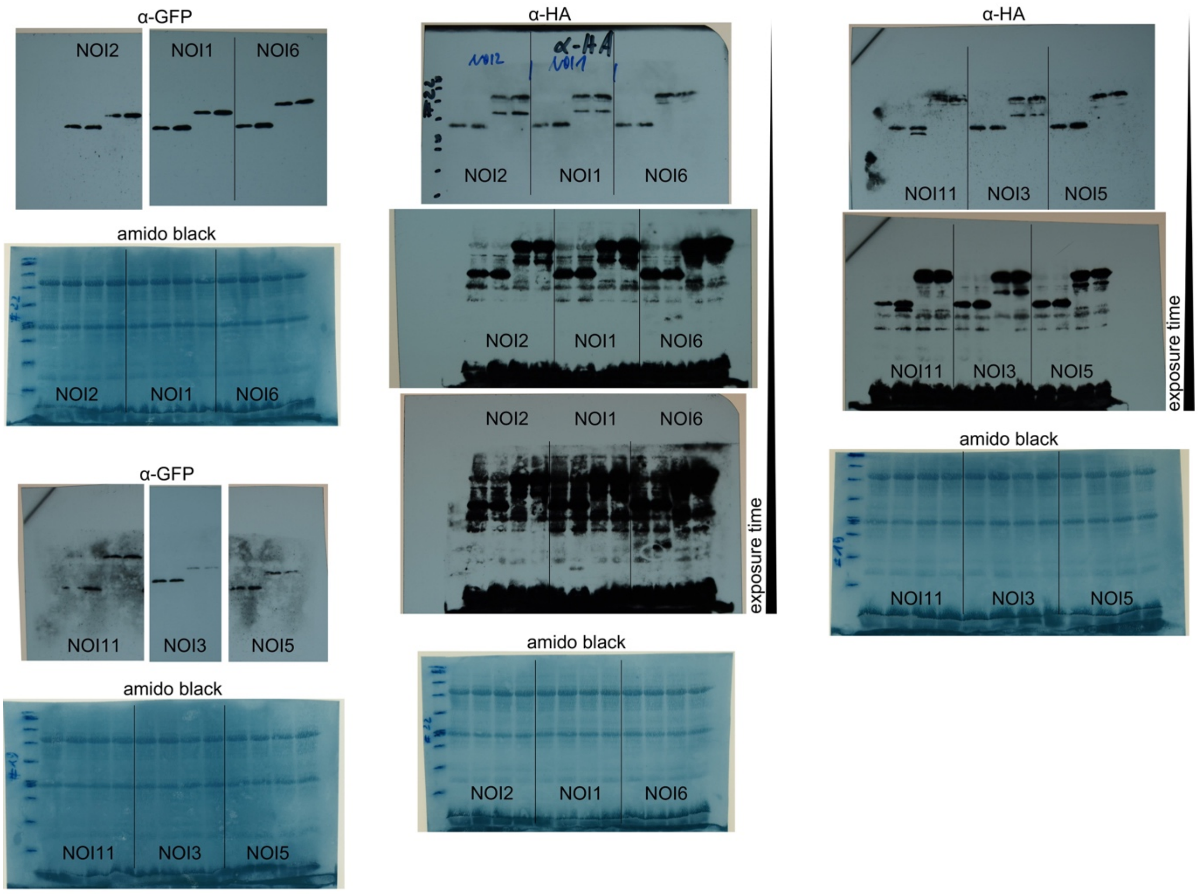


**Figure S7: Uncropped immunoblots shown in Fig. 6.** Amido black-stained membranes and immunoblots are shown. Data is representative of 3 independent replicates

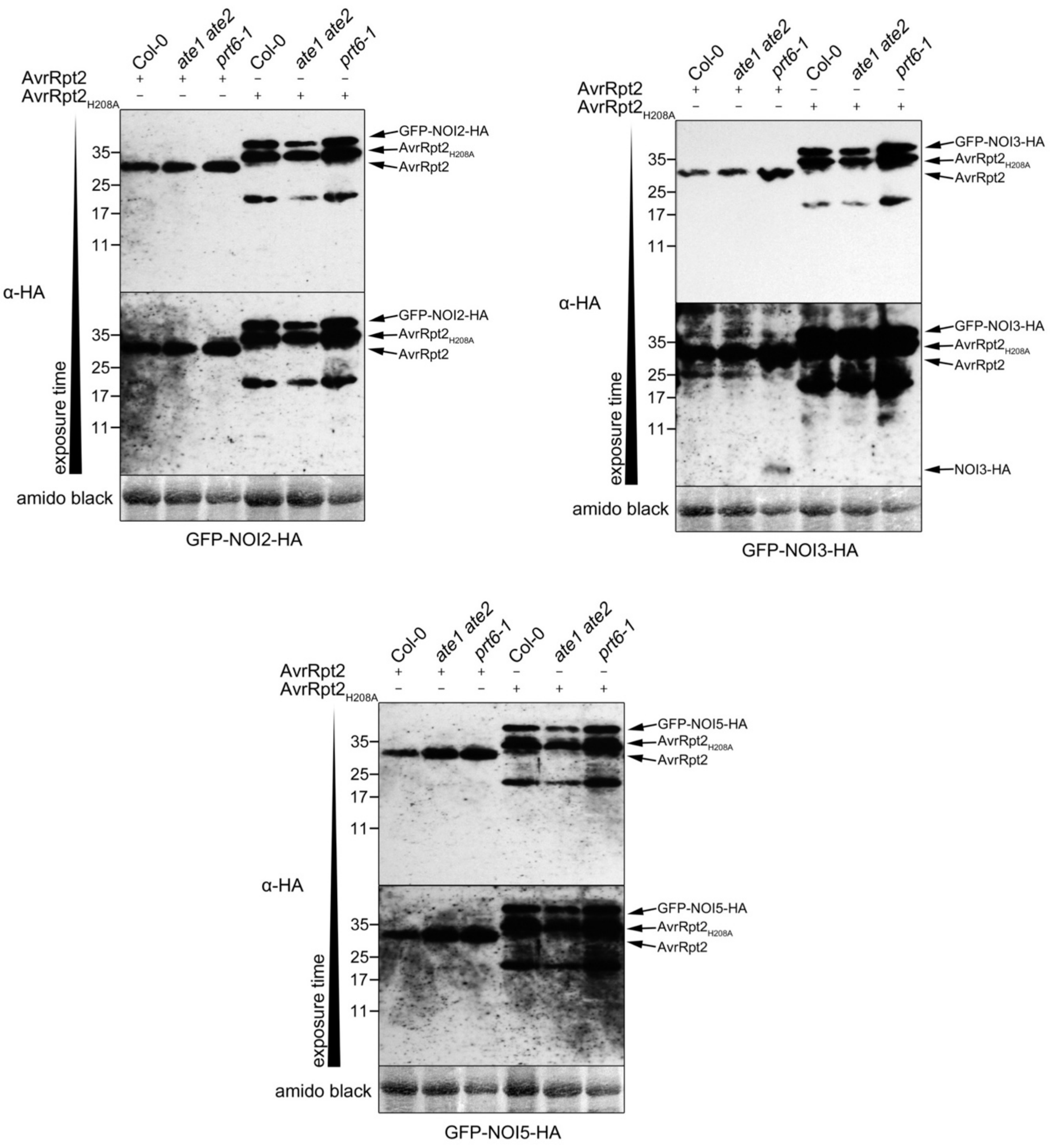


**Figure S8: $^{\Delta}$NOI2-HA, $^{\Delta}$NOI3-HA and $^{\Delta}$NOI5-HA fragment stability in *ate1 ate2* and *prt6-1* mutant protoplasts.** Stability of the wild-type $^{\Delta}$NOI-HA fragments upon transient co-expression of double-tagged NOI2, NOI3 or NOI5 with either wild-type AvrRpt2 or the AvrRpt2$^{H208A}$ inactive variant. Arabidopsis protoplasts derived from wild-type Col-0 plants, as well as *ate1 ate2* and *prt6-1* mutant plants were co-transfected with plasmids coding for the GFP-NOI-HA fusion proteins and with plasmids encoding AvrRpt2 or AvrRpt2$^{H208A}$. $^{\Delta}$NOI-HA C-terminal fragments (indicated with an asterisk) were detected using anti-HA antibodies. Images of the same membrane with increasing exposure times are presented in order to show more clearly the stabilization of the different $^{\Delta}$NOI-HA fragments. For all panels, original data is presented in Supplemental Fig. S9. Data is representative of 3 independent replicates, although in one replicate, expression in the *ate1 ate2* mutant protoplasts was very low.

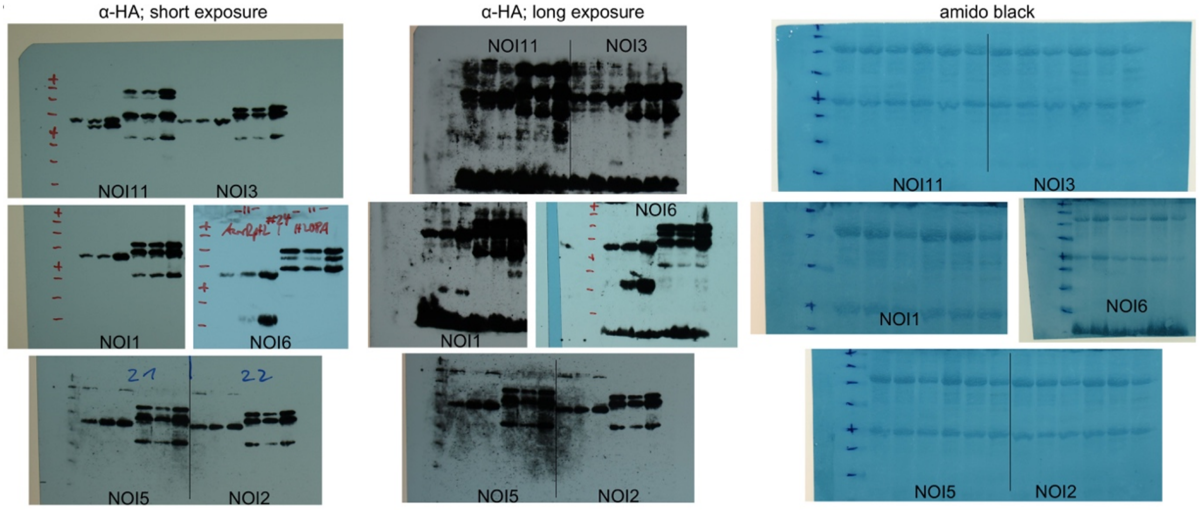


**Figure S9: Uncropped immunoblots shown in Fig. 7.** Amido black-stained membranes and immunoblots are shown. Data is representative of 3 independent replicates, although in one replicate, expression in the *ate1 ate2* mutant protoplasts was very low
